# Characterizing trajectories of innate immune cells in larval zebrafish

**DOI:** 10.1101/2025.08.12.669957

**Authors:** Piyush Amitabh, Raghuveer Parthasarathy

**Affiliations:** Department of Physics, University of Oregon, Eugene, Oregon, USA

## Abstract

It is well established from in vitro studies of immune cells that stimulation by a wide range of potential signals leads to motility and morphology changes. How these physical behaviors manifest inside a living animal remains unclear due to limitations of conventional imaging and analysis approaches. Here, we establish a quantitative framework for imaging and tracking neutrophil and macrophage dynamics in larval zebrafish, spanning a large fraction of the animal for multi-hour timescales with few-minute temporal resolution. We focus especially on the gut, examining innate immune responses to different preparations of the intestinal microbiome. Using light sheet fluorescence microscopy and trajectory analysis of hundreds of individual cells, we characterize speeds, directional persistence measures, and cellular morphology to reveal distinct population behaviors. Individual immune cells exhibit stable motility phenotypes, favoring predominantly motile or non-motile states rather than frequent transitions between them. Gut architecture constrains migration patterns as demonstrated by preferential anterior-posterior movement and a high probability of cells remaining in the vicinity of the gut throughout the imaging duration. Macrophages display significantly reduced sphericity during motile periods compared to non-motile periods, providing a morphological signature that may enable inference of dynamic behavior from static snapshots. Surprisingly, migration patterns remain consistent across diverse microbial conditions – germ-free, conventionally reared, and colonized by two strains of a zebrafish-native *Vibrio* species – indicating that tissue structure exerts a stronger influence than bacterial stimuli on immune surveillance dynamics. Previously observed tissue damage by the wild-type *Vibrio* strain, and the resulting recruitment of immune cells towards the damage site, provided the only microbe-specific cellular behavior. These findings reveal innate immune surveillance as a stereotyped process whose characteristics reflect both cellular decision-making and larger-scale anatomical structure.

## I. INTRODUCTION

The innate immune system serves as the first line of defense against pathogens and supports tissue homeostasis throughout the body [1, 2]. Both functions are especially important around the gastrointestinal tract, which serves as an interface between interior and exterior spaces and between the cells of the host and vast numbers of microbes that include commensal as well as pathogenic organisms. Neutrophils and macrophages, the primary effector cells of innate immunity, communicate with each other and with other cell types through complex, multi-layered chemical networks involving numerous signaling molecules and regulatory pathways [3–6]. Immune cells are also highly motile and physically dynamic, and their strategies for navigating complex, three-dimensional, heterogeneous environments are an integral component of their overall efficacy. In vitro systems, for example collagen gels, have enabled insights into chemotaxis, cell polarity, activation, and force generation by neutrophils and macrophages [7–9]; while they afford experimental control and clarity, their geometry and homogeneity differ considerably from that encountered in live animals. Our understanding of how innate immune cells move in vivo, especially in and around gut-associated tissues, remains minimal, leaving open questions about what cellular features might characterize distinct motility states and whether gut microbes influence surveillance behavior.

Imaging immune cells in live animals is technically challenging but has progressed considerably in recent years. Studies using intravital multi-photon microscopy have revealed the dynamics of immune cells in mammals, but require extensive surgical preparation that can compromise physiological relevance [10–14]. Larval zebrafish (*Danio rerio*), due to their transparency and genetic tractability, provide an especially useful vertebrate model system for studies of the innate immune system [15, 16]. Studies using zebrafish have shown, for example, fewer numbers of gut-associated neutrophils in germ-free animals than in animals colonized by microbes[17, 18], different speeds and modes of migration (mesenchymal versus amoeboid) for neutrophils and macrophages [19], tissue-dependent T-cell motility modes likely reflecting mechanical constraints [20], and a two step search and run response of neutrophils to in-vivo chemical gradients [21]. Furthermore, cutting the caudal fin has become a well-established wound and regeneration assay, which has been used for example to show that neutrophils perform both forward and reverse migration in response to localized damage [22–26], and to test therapeutics for inflammation [27].

Despite the increasing number of investigations, there remains a lack of imaging-based examinations of innate immune cells in volumes large enough to span the entire larval zebrafish gut and over durations large enough to characterize motion types. We therefore used light sheet fluorescence microscopy to image fluorescent neutrophils (transgenic line *Tg(mpx:eGFP)*) and macrophages (*Tg(mpeg1:mCherry)*) [28, 29] in live fish over length scales of millimeters and time scales of a few hours. Light sheet fluorescence microscopy provides rapid, high-volume, three-dimensional imaging with minimal phototoxicity [30–32], and its utility for imaging zebrafish intestinal bacteria and immune cells has been repeatedly demonstrated [33–37]. The resulting image datasets are large, roughly 24 GB per time-point and 1.92 TB per fish for typical dataset, necessitating efficient segmentation and tracking methods to yield immune cell trajectories.

We first validate this imaging and analysis approach with caudal fin cut experiments, providing a test of our analysis methods under conditions of known, directed immune cell migration before applying them to the more complex gut-associated microenvironment. Then we describe neutrophil and macrophage trajectories in larval fish under various gut microbial preparations: germ-free, colonized by ambient microbes (“conventional”), or colonized solely by one of two variants of a zebrafish-isolate *Vibrio* strain, ZWU0020. In past work, we have shown that this strain induces inflammation and other physiological responses in the host animal, responses that require the presence of the actin crosslinking domain (ACD) of the bacterial Type VI Secretion System (T6SS). *Vibrio* with a complete T6SS causes tissue damage, induces macrophage redistribution towards the damage site, and leads to a roughly 100% increase in the amplitude of zebrafish intestinal contractions compared to *Vibrio* lacking the ACD or to non-*Vibrio* bacteria [37]. As described below, we find that under all bacterial conditions, both neutrophils and macrophages show stable motility states, i.e. remaining either motile or non-motile over roughly hour-long durations, maintain proximity to the gut, and migrate preferentially along the intestinal anterior-posterior axis, suggesting that gut architecture is the primary determinant of immune cell movement patterns. Notably, macrophages but not neutrophils show significantly higher sphericity during non-motile periods, providing a morphological indicator of motility state that reflects the greater pseudopodial extension of migrating macrophages.

## II. RESULTS

### A. Validation of 3D immune cell tracking using the caudal fin cut wound model

We performed transverse amputations of the caudal fin in 5 days post-fertilization (dpf) zebrafish larvae, cutting midway between the posterior end of the notochord and the distal tip of the fin (Figure 1A). Using the dual transgenic line *Tg(mpx:GFP; mpeg1:mCherry)*, we simultaneously imaged neutrophils (cells expressing the myeloperoxidase, *mpx*) and macrophages (cells expressing macrophage-expressed gene-1, *mpeg1*) during the wound response [28, 29] (Figure 1B and Supplemental Videos 1, 2). Time-lapse imaging was performed on a custom-built light sheet fluorescence microscope from 1 to 5 hours post-amputation, capturing a tissue volume of approximately 4.8 × 10^7^ *µ*m^3^ (400 *µ*m along the dorsal-ventral, 800 *µ*m along the anterior-posterior, and 150 *µ*m along the left-right axes), with images acquired every 3 minutes.

**FIG. 1:**
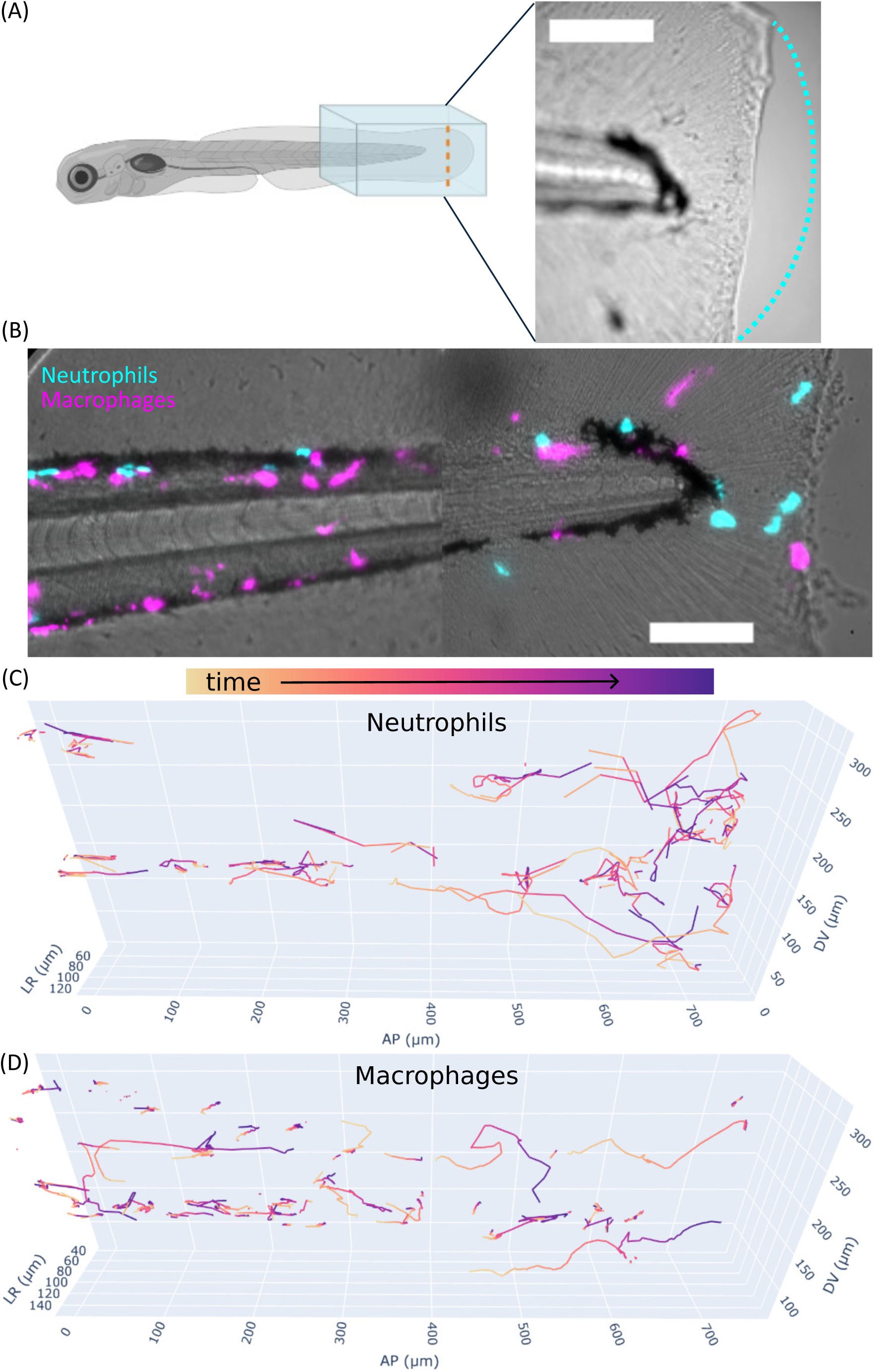
(A) Left: illustration of a larval zebrafish. Blue box indicates the imaging volume (∼800 *µ*m along the anterior-posterior axis), with its posterior boundary aligned to the caudal fin tip; orange dashed line indicates the location of the caudal fin transection. Right: representative brightfield image showing the fin clip, with the original fin shape approximated by the dotted cyan curve. (B) Representative composite image at 1 hour post-fin cut showing brightfield, macrophages (magenta), and neutrophils (cyan). (C, D) Cumulative tracks of neutrophils (C) and macrophages (D) over 1 to 5 hours post-fin cut from a representative specimen. Colors represent normalized time from track initiation (yellow) to termination (purple). Both cell types show migration towards the fin cut site (right). See Supplemental 3D Figures 1–4 for interactive 3D trajectories. AP: anterior-posterior; DV: dorsal-ventral; LR: left-right. All scale bars: 100 *µ*m.

Following image pre-processing and drift correction, individual immune cells were segmented and tracked across time points to generate single cell trajectories (see Materials and Methods). Simple cell counts showed more number of both cell types in the field of view of the fin cut assay than the control (Supplemental Figure 1A). The resulting trajectory data clearly demonstrated directed migration of both neutrophils and macrophages toward the amputation site (Figure 1C, D; Supplemental 3D Figures 1–4), consistent with the known recruitment of immune cells to sites of tissue damage [22–26]. All cell trajectories are provided as Supplemental Data.

**FIG. 2:**
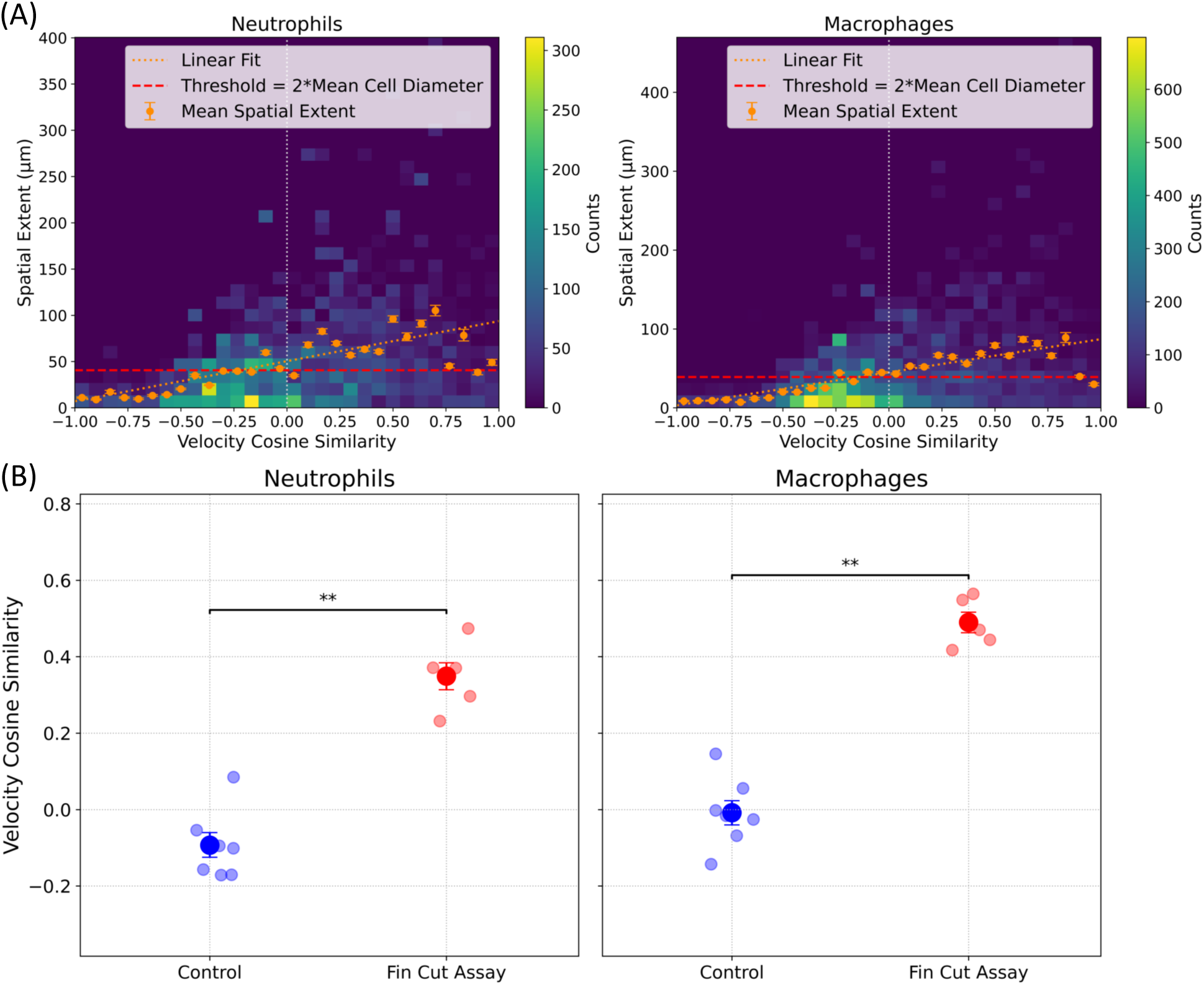
Characterization of immune cell trajectory properties in the caudal fin region for neutrophils (left) and macrophages (right). (A) 2D histograms of spatial extent versus mean velocity cosine similarity for subtracks in the caudal fin region, pooling control and fin-cut fish. The horizontal red line shows the motility classification threshold of twice the mean cell diameter. Orange points show mean spatial extent binned by velocity cosine similarity; the orange dashed line is a linear regression fit to all data points (see main text). (B) Velocity cosine similarity for motile subtracks in fin-cut and control fish. Both cell types show significantly higher velocity cosine similarity for the fin cut assay compared to the control, indicating more consistent directional movement. *n*_control_ = 5, *n*_fin cut assay_ = 7; total *N* = 12. ∗ : *p <* 0.05, ∗∗ : *p <* 0.01.

**FIG. 3:**
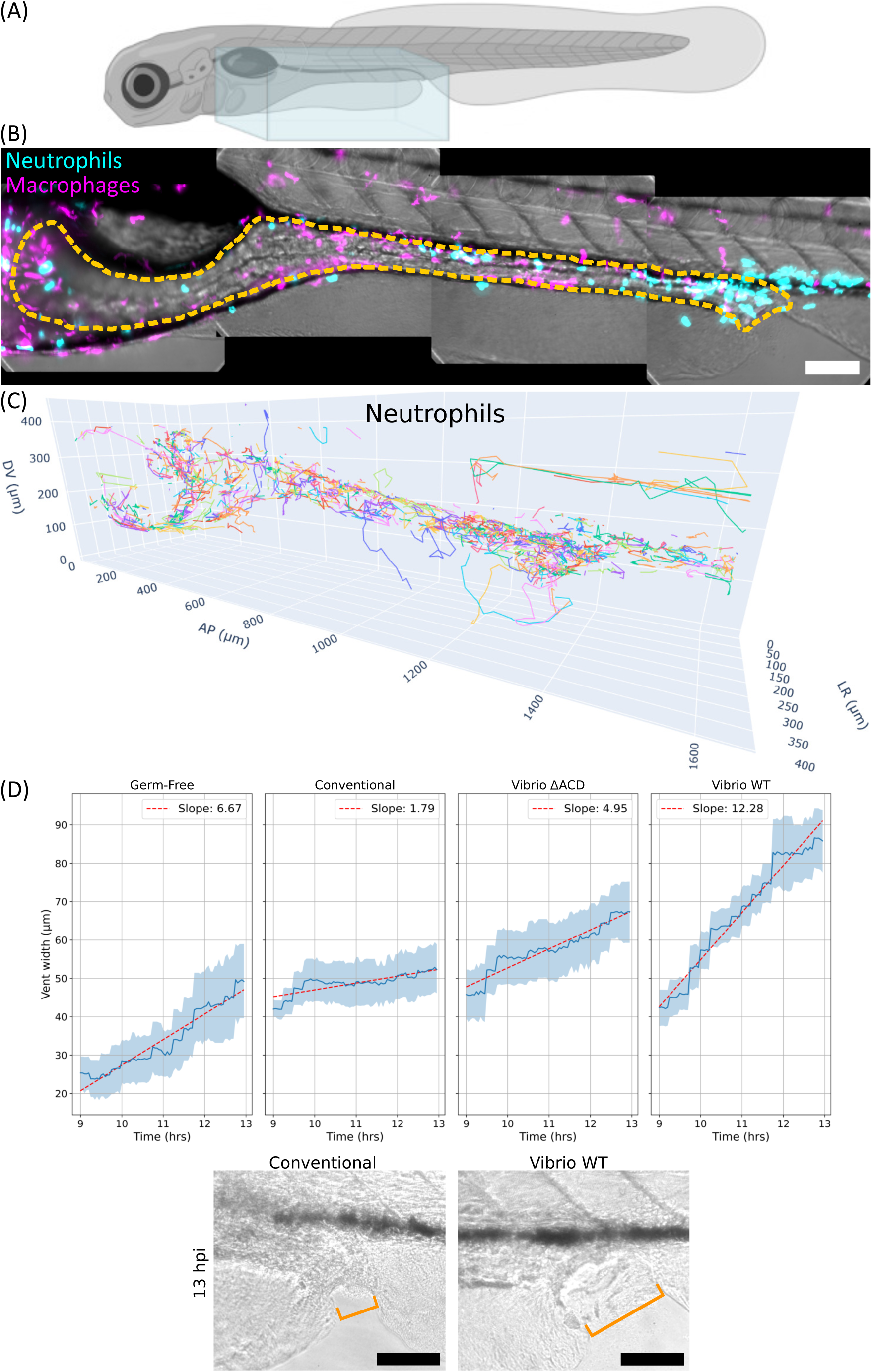
(A) An illustration of a larval zebrafish showing the imaging volume spanning the gut region as a blue box. (B) Composite maximum intensity projection image at a single time point (11 hpi, midpoint of imaging window) showing brightfield (gray), macrophages (magenta), and neutrophils (cyan). The orange dashed line outlines the gut. (C) Cumulative tracks of neutrophils over the 9 to 13 hpi imaging window. Different colors represent individual cell tracks, illustrating the dynamic movement of neutrophils. (D) Upper: Zebrafish vent width over the course of imaging for the different experimental groups: Germ-free, conventionally reared (Conventional), *Vibrio*^Δ*ACD*^-inoculated, and *Vibrio^W^ ^T^* -inoculated fish. Increased vent width is an indicator of tissue damage. *n*_Germ Free_ = 5, *n*_Conventional_ = 5, *n*_Vibrio ΔACD_ = 5, *n*_Vibrio WT_ = 6; total *N* = 21. Lower: Brightfield images showing vent widths for conventionally raised and *Vibrio^W^ ^T^* -inoculated fish at the end of the imaging window. Orange lines indicate the vent width. All scale bars: 100 *µ*m.

**FIG. 4:**
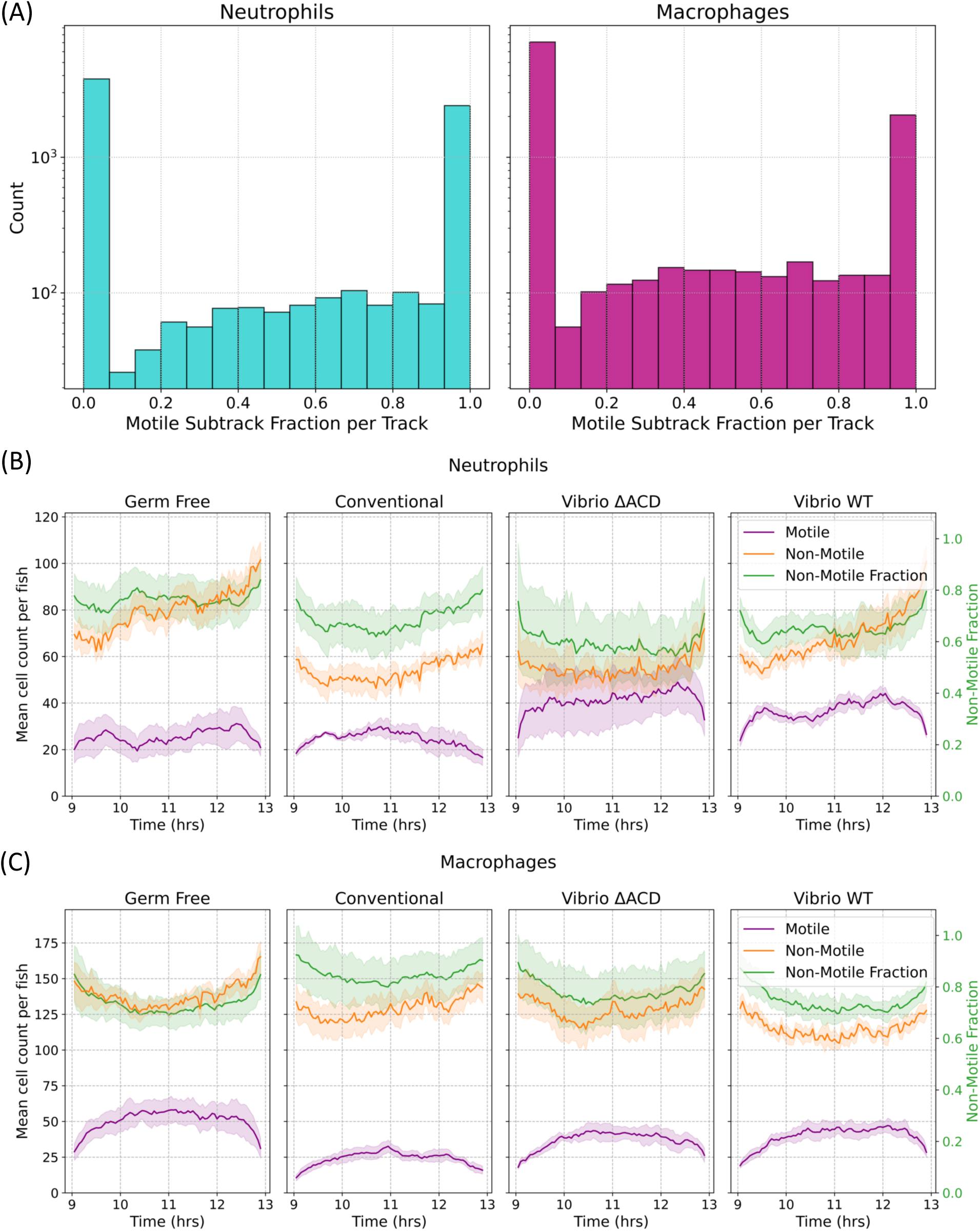
(A) Histograms showing the motile subtrack fraction per track for neutrophils (left) and macrophages (right), pooled across all experimental groups. The distribution shows peaks at 0 and 1, indicating that immune cells generally adopt a consistent motile or non-motile phenotype over the 4-hour observation period, though a subset of cells exhibits intermediate behavior. Bin width = 0.06. (B, C) Counts of motile and non-motile subtracks, and the ratio of non-motile to total subtracks from 9 to 13 hpi for (B) neutrophils and (C) macrophages across experimental groups: Germ-free, conventionally reared (Conventional), *Vibrio*^Δ*ACD*^-inoculated, and *Vibrio^W^ ^T^* -inoculated fish. Motile cell numbers remain relatively constant across conditions. Error bars represent uncertainties from bootstrapping. *n*_Germ Free_ = 5, *n*_Conventional_ = 5, *n*_Vibrio_ ΔACD = 5, *n*_Vibrio_ WT = 6; total *N* = 21.

### B. Establishment of motion classification criteria

To extract meaningful physical parameters from cellular trajectories, we developed a computational framework that accounts for the intermittent nature of immune cell migration. Individual cells exhibit alternating periods of active migration and relative quiescence, a characteristic feature of immune cell behavior in tissue environments [38]. To capture this temporal heterogeneity, we subdivided each complete trajectory into “subtracks” based on frame-to-frame displacement. Specifically, a new subtrack was initiated whenever the displacement transitioned above or below a threshold of half the mean cell diameter per 3 minute interval. We computed the spatial extent of each subtrack as the diameter of the smallest circumscribing sphere. We calculated the velocity cosine similarity of each subtrack, a measure of directional persistence defined as the cosine of the angle between consecutive velocity vectors, averaged over the subtrack. Evaluating immune cells in the caudal fin area, pooling both cut and control fish, plots of spatial extent versus velocity cosine similarity for each subtrack show a positive correlation for both neutrophils and macrophages (Figure 2A), indicating that more spatially extended tracks are also more directionally persistent.

To compare this correlation with what would arise from undirected random motion with the same track features and sampling as in the data, we generated simulated tracks by randomizing step directions while preserving each track’s step-size distribution and track length, with detector noise matching experimental imaging conditions (see Materials and Methods). Simulated tracks showed no such correlation, with velocity cosine similarity clustered near zero regardless of spatial extent (Supplementary Figure 11A), and both velocity cosine similarity and velocity anisotropy being significantly lower than experimental values (Supplementary Figure 11B, C). These results confirm that the observed directional signatures reflect intrinsic directional bias rather than kinematic artifacts. To classify subtracks as motile or non-motile, we define a simple threshold of spatial extent being at least twice the mean cell diameter. To justify this beyond its intuitive validity, we consider for each cell type a linear fit of spatial extent versus velocity cosine similarity; the spatial extent corresponding to zero velocity cosine similarity is 47.2 *µ*m (*R*^2^ = 0.109*, n* = 13882 datapoints) for neutrophils and 42.6 *µ*m (*R*^2^ = 0.115*, n* = 23765) for macrophages, close to twice the mean cell diameter of 20.4 *µ*m and 19.5 *µ*m for neutrophils and macrophages respectively (Figure 2A). Since zero velocity cosine similarity corresponds to random or repetitive motion, including physiological noise from blood flow and small displacements from cell shape changes, this agreement implies that our motility threshold effectively separates persistent migration from non-directional motion.

Applying this motility classification scheme to the fin cut assay demonstrates its effectiveness in capturing differences in immune cell behavior. Motile subtracks from fin-cut fish exhibited significantly higher velocity cosine similarity compared to motile subtracks from the uninjured control group (Figure 2B), confirming that our analysis pipeline successfully identified the expected increase in directional migration of both neutrophils and macrophages following tissue damage. In control fish, motile subtracks show velocity cosine similarity near zero, indicating largely random motion; small deviations from zero reflect residual noise not fully eliminated by the analysis pipeline. In injured fish, the innate immune cells show persistent motion, visually apparent as directed toward the wound site.

Having validated our motion analysis pipeline, we turn to investigations of immune cell behavior in response to different gut microbiota inoculations.

### C. Three-dimensional immune cell tracking reveals gut-associated motility patterns

To investigate immune cell dynamics around the larval zebrafish gut in response to varied commensal bacterial groups, we assessed germ-free fish, conventionally reared fish with normal parental tank microbiota, and initially germ-free fish inoculated at 5 or 6 days postfertilization (dpf) with one of two variants of the zebrafish-native *Vibrio* strain ZWU0020. As noted above, we showed in earlier work that colonization with wild-type *Vibrio* ZWU0020 (which we denote *Vibrio^W^ ^T^*) causes tissue damage near the zebrafish vent (the posterior opening of the gut), which recruits macrophages to the vent from their normal midgut positions during a period of roughly 14 to 20 hours post inoculation (hpi) [37]. Vent damage and immune activation are minimal, with levels similar to germ-free fish, in larvae colonized with a *Vibrio* mutant lacking the actin crosslinking domain (ACD) of the Type VI Secretion System (T6SS), (denoted *Vibrio*^Δ*ACD*^) [37]. In prior work [37], macrophages positions were assessed at 20 minute intervals and neutrophils were not tracked.

In the studies described here, we began imaging at 9 hpi and tracked neutrophil and macrophage cells at 3-minute intervals through 13 hpi. This time window was chosen to investigate and capture immune cell behavior prior to the major macrophage redistribution to the vent previously observed at 14-20 hpi [37]. Gut dissection and plating at the end of the imaging period (13 hpi) confirmed successful *Vibrio^W^ ^T^* colonization at 10^3^ to 10^4^ colony forming units (CFU) per gut (see Materials and Methods). Imaging of *Vibrio^W^ ^T^* -dTomato fluorescence at representative timepoints (10, 11, and 13 hpi) showed bacterial localization primarily at the gut bulb and the vent (see Materials and Methods; Supplemental Figure 2 and Supplemental Video 3). Germ-free, conventionally reared, or *Vibrio*-inoculated zebrafish were imaged with light sheet fluorescence microscopy at 3-minute intervals at 5 or 6 dpf from 9 to 13 hpi for inoculated fish or the same age for other non-inoculated fish. The imaging volume covered the entire gut: approximately 1600 *µ*m along the anterior-posterior (AP) axis, 100 *µ*m left-right (LR), and 400 *µ*m dorsal-ventral (DV) (Figure 3A). Using the dual transgenic line *Tg(mpx:GFP; mpeg1:mCherry)*, we imaged and tracked individual neutrophils and macrophages throughout the 4-hour observation period (Figure 3B, C and Supplemental Video 4, 5). All cell trajectories are provided as Supplementary Data. The mean track lengths were 24 and 32 frames (69 and 93 minutes) for neutrophils and macrophages, respectively,, pooled across all groups. These cells showed a similar relationship between spatial extent and velocity cosine similarity as immune cells in the fin cut assay (Supplemental Figure 3). As with the fin cut data, simulated random walks showed no such correlation, with velocity cosine similarity clustering around zero, while experimental data showed significantly higher directional persistence across nearly all conditions (Supplemental Figure 12A, B). Vent width measurements confirmed the expected T6SS-dependent tissue damage at the end of the imaging window, with *Vibrio^W^ ^T^* -associated fish showing the largest increase compared to other groups (Figure 3D, Supplemental Figure 4).

### D. Immune cells maintain distinct motility states with robust motility patterns across microbial conditions

Using our framework for classifying the motility of subtracks, we found that both neutrophils and macrophages showed a tendency to remain either motile or non-motile for extended periods of time comparable to the mean track duration of over one hour rather than showing frequent transitions between motility states. Histograms of the motile sub-track fraction per track, grouped across all bacterial conditions, reveal bimodal distributions with peaks at 0 and 1 for both neutrophils and macrophages (Figure 4A; bin width 0.06). This bimodal pattern contrasts with the uniform distribution expected if there were frequent switching between motility states. While a subset of cells exhibits intermediate fractions of motile behavior, the observed distribution indicates that innate immune cells preferentially adopt and maintain a consistent motile or non-motile phenotype over the four-hour observation period. Analysis included only tracks with minimum track lengths of 5 frames to ensure robust motility classification. This distribution pattern was consistent across all four experimental groups and in the fin cut assay (Supplemental Figure 5, 6).

To more finely assess motility behaviors and possible dependence on microbial stimulation, we examined the number and fraction of motile and non-motile cells (subtracks) at each timepoint from 9 to 13 hpi. The numbers of motile neutrophils and macrophages remained fairly constant and similar across all four conditions (germ-free, conventional, *Vibrio^W^ ^T^*, and *Vibrio*^Δ*ACD*^) (Figure 4B, C). Non-motile neutrophils increased in number over time in *Vibrio^W^ ^T^*-inoculated fish (Figure 4B), which was mainly driven by the increase in the number of non-motile neutrophils near the vent region (Supplemental Figure 7). Macrophages showed no significant temporal changes in non-motile subtrack numbers across any group (Figure 4C). We note that for some experimental conditions there is an initial increase and a late decrease in motile cells (Fig. (Figure 4B,C). This is a likely consequence of the requirement of a minimum 5 frame (15 minute) track length, as motile cells have an appreciable likelihood to enter or leave the field of view during the peripheral 15 minutes, and so are not counted.

### E. Macrophage morphology correlates with motility state

To investigate whether immune cell motility correlates with detectable morphological changes, we quantified cellular sphericity, defined as the ratio of the surface area of a sphere with the same volume as the measured cell to the measured surface area of the cell. Perfect spheres have a sphericity value of 1.0, whereas elongation or any other shape distortions reduce this value.

Neutrophils exhibited no significant differences in sphericity between motile and non-motile states across all experimental groups (Figure 5A). In striking contrast, macrophages demonstrated significantly higher sphericity during non-motile periods compared to motile periods, indicating increased cellular rounding when stationary. Representative microscopy images reveal that motile macrophages display prominent pseudopodial protrusions, whereas non-motile macrophages adopt a more rounded morphology (Figure 5B and Supplemental Videos 6– 9). This morphological distinction suggests that motile macrophages extend pseudopodia more extensively than their non-motile counterparts, providing a snapshot-based indicator of motility state that does not require temporal tracking to assess. This behavior is also observed in the response of these cells in the fin cut assay (Supplemental Figure 1B). We examined numerous additional cellular parameters, including volume and normalized anterior-posterior position, but sphericity was the only metric that distinguished motile from non-motile behavior across all macrophage groups (Supplemental Figure 8A, B and 1C).

**FIG. 5:**
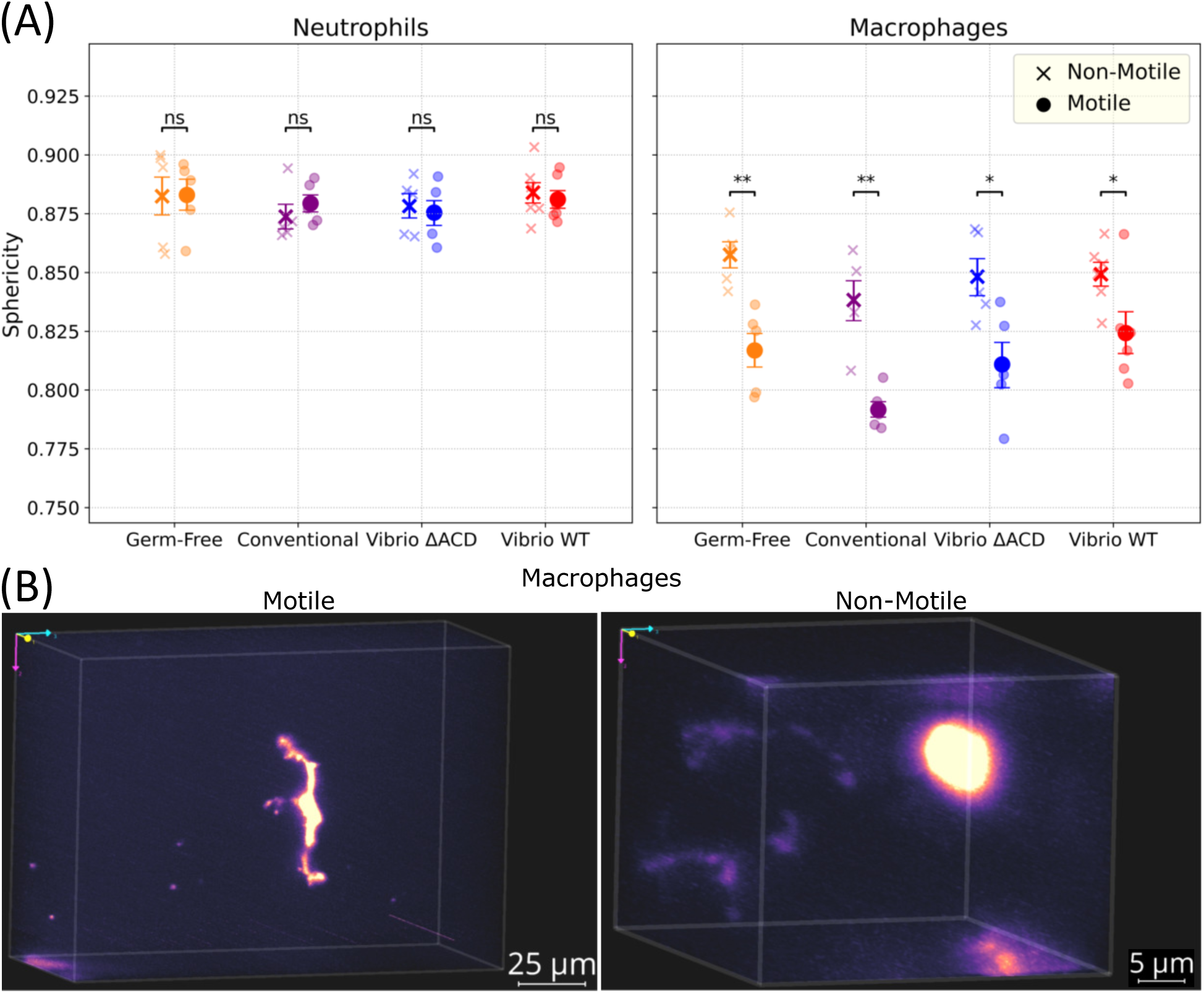
Macrophages exhibit reduced sphericity associated with pseudopodial extensions during motile phases. (A) Sphericity comparison between motile and non-motile subtracks for neutrophils (left) and macrophages (right) across all experimental groups. Macrophages exhibit significantly lower sphericity during motile periods compared to non-motile periods, while neutrophils show no significant difference between motility states. (B) Representative images of macrophages during motile (left) and non-motile (right) phases. Motile macrophages display prominent pseudopodial protrusions, whereas non-motile macrophages adopt a more rounded morphology (Supplemental Videos 6– 9). Error bars represent uncertainties from bootstrapping. *n*_Germ Free_ = 5, *n*_Conventional_ = 5, *n*_Vibrio ΔACD_ = 5, *n*_Vibrio WT_ = 6; total *N* = 21. ns : not significant, ∗ : *p <* 0.05, ∗∗ : *p <* 0.01.

### F. Gut architecture constrains immune cell migration patterns

We manually segmented images to identify gut-associated regions, including the intestinal lumen and surrounding tissue (Figure 3B and Supplemental Video 10). We classify as “gut-associated” immune cells that are located within these boundaries at any point during imaging. To assess the persistence of gut association over extended periods, we considered subtracks with a duration of at least five frames and examined the fraction of time spent in the gut volume. Non-motile subtracks spent more than 80% of their time within the gut, considering averages across all experimental groups(Figure 6A). Surprisingly, motile subtracks also demonstrated high gut residence probability (*>* 70% across all groups). Simulating random walks with the same average frame-to-frame displacement as immune cells and assessing their residence in a volume like that of the larval gut gives a residence probability of 30−40% for walkers with the same median track lengths of the observed neutrophils and macrophages, indicating that gut-associated immune cells migrate predominantly along the boundary of the gut.

**FIG. 6:**
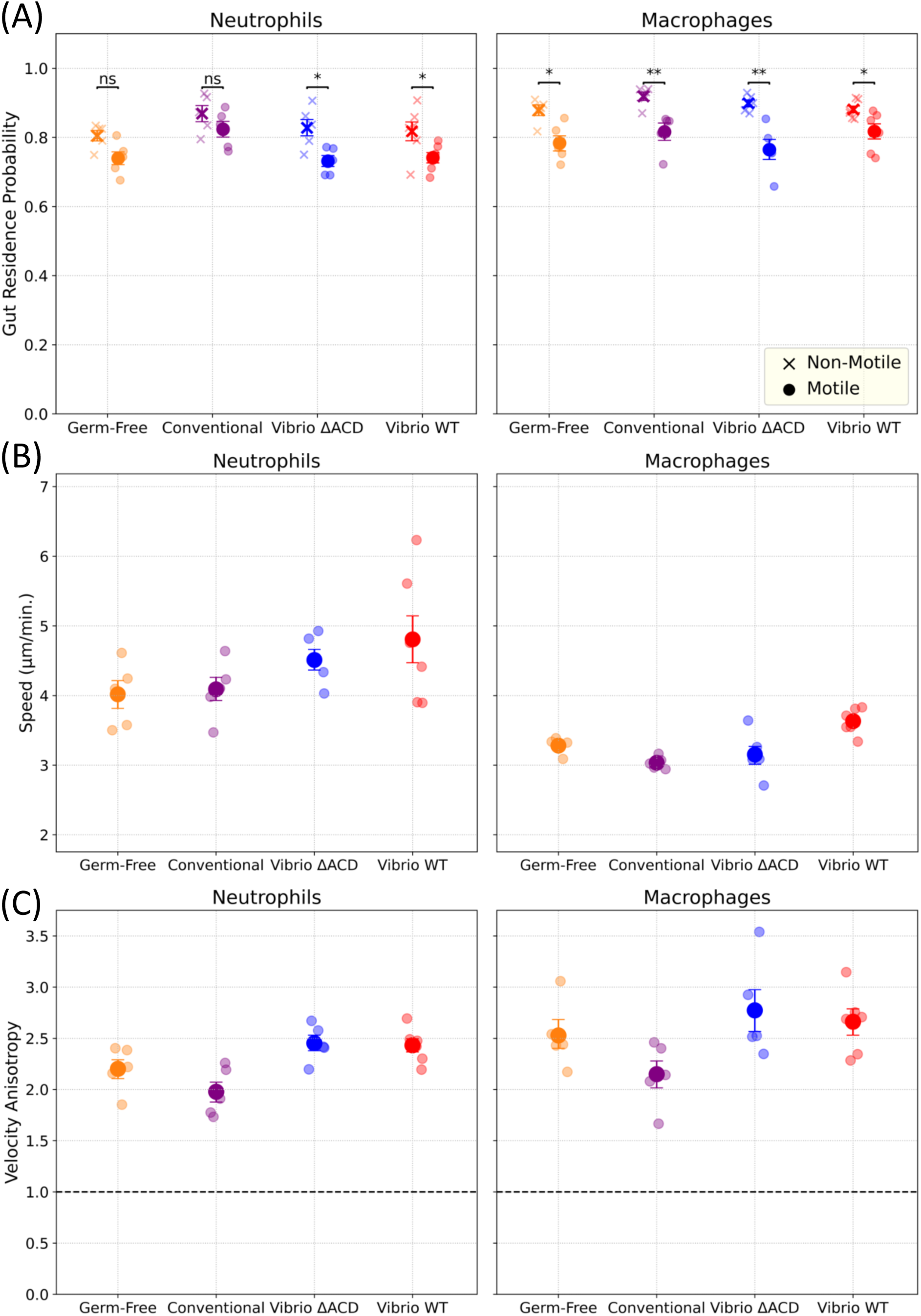
Characterizations of immune cell residence and migration dynamics in gut-associated regions for neutrophils (left) and macrophages (right). (A) Gut residence probability, defined as the fraction of time a cell that is ever in the vicinity of the gut remains in the vicinity of the gut, for motile and non-motile subtracks, demonstrating that both populations spend the majority of their time within gut-associated regions across all experimental conditions. (B) Average speed of motile subtracks, calculated from frame-to-frame displacements, shows no significant differences between experimental groups for either cell type. (C) Velocity anisotropy quantifies preferential movement along the anterior-posterior axis relative to perpendicular directions. Values consistently exceed 1.0 across all conditions, indicating migration primarily along the intestine rather than in dorsal-ventral or left-right directions. Error bars represent uncertainties from bootstrapping. *n*_Germ Free_ = 5, *n*_Conventional_ = 5, *n*_Vibrio ΔACD_ = 5, *n*_Vibrio WT_ = 6; total *N* = 21. ns : not significant, ∗ : *p <* 0.05, ∗∗ : *p <* 0.01.

Analysis of motile subtrack characteristics revealed that most kinetic parameters, including mean speed and velocity components, showed no significant differences between experimental groups (Figure 6B and Supplemental Figure 9). We also provide these parameters for the fin cut assay (Supplemental Figure 10). However, we found that immune cell migration shows a strong directional bias. We calculated all cells’ velocity anisotropy, given by:

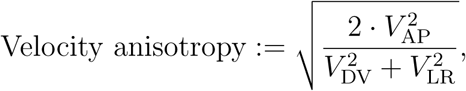

where *V*_AP_, *V*_DV_, and *V*_LR_ represent velocity components along the anterior-posterior, dorsal-ventral, and left-right axes, respectively. Velocity anisotropy consistently exceeded 1.0, the value for isotropic motion, across all experimental conditions (Figure 6C), and was significantly higher than the anisotropy of simulated random walks drawn from the experimental distribution of step sizes (see Materials and Methods; Supplemental Figure 12C), indicating preferential movement along the anterior-posterior axis compared to dorsal-ventral or left-right directions.

Combined with the high gut residence probability of motile cells, this result demonstrates that gut-associated immune cells migrate primarily along the intestinal length (along anterior-posterior axes) rather than in left-right or dorsal-ventral directions, suggesting that gut architecture fundamentally constrains immune cell movement patterns regardless of microbial stimulus.

## III. DISCUSSION

To date, there have been few quantitative analyses of immune cell dynamics in living organisms over volumes that span a large fraction of the animal, and to our knowledge, this study is the first that characterizes innate immune dynamics around the gastrointestinal tract. Given strong, multifaceted interactions between the intestinal microbiota and the host immune system [39–42], it is natural to expect that different microbial stimuli should lead to different immune cell motility patterns. Surprisingly, we find little difference in motility characteristics in germ-free larval zebrafish, conventionally reared fish, and fish inoculated with a pathobiont *Vibrio* species, other than spatial redistribution to the site of *Vibrio*-induced tissue damage [37]. Immune cell motion is, however, strongly influenced by the anatomical context of the host. Neutrophils and macrophages that encounter the gut tend to spend a high fraction of their time near the gut, and migrate largely along the anterior-posterior axis. This strong axial preference was consistent across all microbial conditions, indicating that gut structure, not microbial signals, primarily dictates migration patterns.

The remarkable consistency of immune cell behavior across microbial conditions – germ-free, conventional, *Vibrio^W^ ^T^*, and *Vibrio*^Δ*ACD*^ – demonstrates robust migration patterns independent of bacterial stimuli. The bimodal distribution of motility states, constant numbers of motile cells, and invariant kinetic parameters across all conditions suggest that gut architecture exerts stronger influence than microbial signals during the 9-13 hpi window. Only the localized accumulation of non-motile neutrophils near *Vibrio*-induced tissue damage deviated from this pattern, indicating that while bacterial activity can create specific retention sites, it does not fundamentally alter migration dynamics. These findings suggest that therapeutic strategies targeting immune motion should consider the dominant role of tissue architecture over microbial influences.

On the timescale of a few hours, we found stable motility states (either motile or non-motile) for both neutrophils and macrophages, rather than frequent transitions in behavior. While our study made use of live imaging over several hours, it is interesting to ask whether motility states can be gleaned from single snapshots, potentially even from non-living animals or histological samples. We found no notable morphological differences between motile and non-motile neutrophils. Macrophages, in contrast, were much less spherical when motile. This feature may allow the inference of macrophage dynamic behavior from static snapshots alone. Due to the lack of cell membrane and actin markers, we cannot definitively classify the cellular protrusions observed in the motile and less spherical subset of macrophages, though visual inspection suggests these protrusions are consistent with the general form of pseudopodia (see Supplementary Videos 6– 9). Cellular protrusions can be classified into several types based on their morphology and cytoskeletal architecture, like lamellipodia (broad, sheet-like projections), filopodia (thin, finger-like extensions), and blebs (pressure-driven spherical extensions) [43]. Characterizing the protrusion types present in migrating gut macrophages, along with their mechanical characteristics and modes of migration (amoeboid versus mesenchymal), represents an interesting avenue for future investigation.

A key open question concerns the mechanistic basis of immune cell guidance in vivo. Neutrophils and macrophages are well known to migrate along chemokine gradients toward sites of infection or tissue damage [5], yet the striking anatomical constraint we observe, cells remaining in the vicinity of the gut and migrating along its axis, suggests that physical cues also play a role. The gut is a tubular structure, presenting curved surfaces that could engage curvotaxis, whereby cells preferentially migrate along regions of negative curvature through nucleus-mediated mechanosensing [44, 45]. Cell migration can also be directed by durotaxis (stiffness gradients) and haptotaxis (substrate-bound signals) [46, 47]. The extracellular matrix in zebrafish undergoes substantial remodeling during development, with region-specific changes in composition and mechanical properties [48, 49]. Given that different anatomical regions exhibit distinct ECM characteristics at different developmental stages, the relative contributions of chemotaxis, curvotaxis, and durotaxis to immune cell navigation likely vary across tissue contexts. Future work employing membrane and cytoskeletal markers, combined with ECM characterization, would help dissect these contributions.

Our study focused on neutrophils and macrophages in larval zebrafish at 5–6 days post-fertilization, a stage when innate immunity predominates and adaptive immune cells have not yet matured [50]. Our study focused on neutrophils and macrophages in larval zebrafish at 5–6 days post-fertilization, a stage when innate immunity predominates and adaptive immune cells have not yet matured [50]. Notably, heterogeneous motility phenotypes within immune cell populations have been observed in other contexts. In lymph nodes, T cell populations contain both highly motile naive cells undergoing random walks and arrested cells forming stable conjugates with dendritic cells during antigen recognition, with cells maintaining their respective motility states for periods of 8–12 hours [13, 14, 51]. Similarly, heterogeneity in migration modes — mesenchymal versus amoeboid — has been described for neutrophils and macrophages in zebrafish tissues [19]. Whether the stable motile and non-motile phenotypes we observe in gut-associated immune cells correspond to distinct functional subtypes, analogous to the naive versus antigen-engaged distinction in T cells, or represent different stages of a behavioral program, remains an open question. More generally, motility patterns across the range of immune cell types remain under-explored. The immune systems of zebrafish and mammals are highly conserved but differ in key aspects, including, in zebrafish, the absence of lymph nodes and reliance on the kidney as the primary hematopoietic organ [52, 53]. Extending our framework to mammals with more complex organ structure and tissue compartmentalization, to later developmental stages when adaptive immunity is established, and to other immune cell types would be important next steps and would reveal whether the behaviors uncovered in innate immune cells in larval zebrafish are reflected in other organisms and other stages of development. The ability to track hundreds of cells in 3D across multi-hour timescales, combined with emerging single-cell transcriptomic atlases [54], could ultimately link motility phenotypes to transcriptional states.

The combination of stable motility states, architectural constraints, and cell-type-specific morphological signatures reveals immune surveillance as a stereotyped process whose characteristics reflect both cellular decision-making and larger-scale structure. The quantitative framework established here and the open-source computational pipeline we provide should allow translation to other model systems and cell types, identifying conserved versus context-specific features of immune surveillance mechanisms during development and disease.

## IV. MATERIALS AND METHODS

### A. Zebrafish husbandry and transgenic lines

All experiments with zebrafish followed standard procedures [55] and were performed in accordance with protocols approved by the University of Oregon Institutional Animal Care and Use Committee (protocol numbers AUP-22-02 and AUP-20-16). Fish from which the larvae used in these studies were derived were maintained in the University of Oregon Zebrafish Facility at 28°C with a 14-h/10-h light/dark cycle.

The transgenic line Tg(mpeg1:mCherry), allele number: gl23, is described in reference [28]. The transgenic line Tg(tnf*α*:GFP);Tg(mpeg1:mCherry), allele number pd1028Tg, is described in reference [56].

### B. Gnotobiotic derivation and bacterial colonization

Embryos were derived germ-free following established protocols [57]. Fertilized eggs were incubated for 6 hours in sterile embryo medium (EM) containing antibiotics (100 *µ*g/mL ampicillin, 10 *µ*g/mL gentamycin, 1 *µ*g/mL tetracycline, 1 *µ*g/mL chloramphenicol, 250 ng/mL amphotericin B). After surface sterilization with 0.003% sodium hypochlorite and 0.1% polyvinylpyrrolidone-iodine, embryos were distributed into T25 flasks at one embryo per mL in 15 mL sterile EM.

*Vibrio* ZWU0020 (PRJNA205585) was previously isolated from the zebrafish intestinal tract. From freezer stocks kept at -80°C, cultures were grown overnight in Luria Broth at 30°C with agitation. Then, bacterial cultures were pelleted through centrifugation for 3 min at 7,000 × g, washed once in sterile EM, and inoculated at 10^6^ CFU/mL directly into flask water. Larvae were colonized at 5 or 6 dpf, with time-lapse imaging from 9 to 13 hpi.

### C. Construction of *Vibrio* ZWU0020 T6SS mutants

The actin crosslinking domain deletion mutant (Vibrio^ΔACD^) was constructed using markerless allelic exchange with the pAX2 vector as previously described [37, 58]. In brief, splice-by-overlap extension generated deletion cassettes with the following primers: 5’ HR: 5’-CCCAATGATAGCCACGGTTG-3’ + 5’-CCATTCCATTTTCCACTAGGCTAAAGGACACACCTT-3’; 3’ HR: 5’-AAGGTGTGTCCTTTAGCCTAGTGGAAAATGGAATGG-3’ + 5’-GGCGCAAGATTTTCAATCA-3’. After conjugation into *Vibrio* using *E. coli* SM10, merodiploids were selected on TSA-gentamicin at 37°C. Deletion was confirmed by PCR using flanking primers: 5’-CGGAGCTTTGGTCAATCTCA-3’ + 5’-AGGTCTCTCCGTGGAAAACA-3’.

### D. Caudal fin amputation

For wound response assays, 5 dpf larvae were anesthetized in 120 *µ*g/mL MS-222 before mounting on glass slides using 4% methyl-cellulose. The caudal fin was transected midway between the notochord end and fin tip using a sterile scalpel. Larvae recovered in fresh EM at 28^◦^C before imaging.

### E. Quantification of intestinal bacterial abundance

To quantify intestinal bacterial abundance, larval zebrafish were euthanized by hypothermic shock at the indicated timepoints (hours post-inoculation, hpi). Intestines were removed under a dissection microscope, placed in 1 mL sterile embryo medium, and homogenized with zirconium oxide beads using a Bullet Blender^®^ tissue homogenizer (Next Advance, Troy, NY). Homogenates were serially diluted 10^−1^ and 10^−2^, and 100 *µ*L volumes were spread onto LB-agar plates. Plates were incubated at 30^◦^C overnight, and colony-forming units (CFU) were enumerated to calculate bacterial load per intestine.

### F. Light sheet fluorescence microscopy

Imaging was performed using a custom-built light sheet fluorescence microscope described in detail elsewhere [59, 60]. A galvanometer-scanned light sheet of excitation illumination from 448 or 561 nm lasers (Coherent Sapphire, operating at 5 mW) was oriented perpendicular to the optical axis of a 40×, 1.0 NA water-immersion detection objective. Images were captured by a 5.5 MP sCMOS camera (Excelitas pco.edge). Three-dimensional volumes were acquired by scanning the sample along the detection axis in 1 *µ*m steps with 33 ms exposures per plane. The resulting image resolution is approximately 0.4 *µ*m in the dimensions parallel to the sheet and 5 *µ*m perpendicular to the sheet. All hardware components were controlled using *µ*Manager software [61].

For gut imaging, four overlapping sub-regions covering the entire intestine (∼ 1,200 × 300 × 150 *µ*m^3^) were sequentially acquired and computationally registered. Time series were collected at 3-minute intervals for 4 hours (9-13 hpi). For fin wound assays, single volumes were acquired at 3-minute intervals for 4 hours (1-5 hours post-amputation). Original voxel dimensions were 1 *µ*m (z-axis) × 0.1625 *µ*m (y-axis) × 0.1625 *µ*m (x-axis).

### G. Sample mounting

Larvae were anesthetized in 120 *µ*g/mL MS-222 and embedded in 0.7-1.0% low-melt agarose within glass capillaries. Specimens were partially extruded to avoid optical interfaces and oriented with ventral illumination. Up to four samples were mounted simultaneously in sterile EM at 28°C.

### H. Bacterial distribution analysis

To quantify bacterial distribution along the gut during the imaging period, we analyzed *Vibrio^W^ ^T^* -dTomato fluorescence. To reduce host cell and background autofluorescence interference, we created 3D gut lumen masks to restrict analysis to regions where bacteria are present in the zebrafish gut. Using bright-field and dTomato fluorescence images, 2D gut lumen boundaries were manually drawn at every 10th z-plane for each field of view. These 2D masks were extruded across the intermediate planes to generate complete 3D volumes, then stitched across all imaging positions to cover the entire gut (Supplemental Video 3). The gut was masked using this 3D mask and the resulting image was median background subtracted. Pixel intensities were binned along the anterior-posterior axis to generate line profiles of bacterial distribution. Anterior-posterior position is normalized such that 0 and 1 represent the start and end of the gut, respectively.

### I. Gut segmentation

Binary masks that identified the larval zebrafish gut region in images were manually segmented using a Wacom Cintiq^®^ graphics tablet on brightfield images acquired at the median sagittal plane. To optimize efficiency while maintaining accuracy, masks were initially drawn at every 10th timepoint and subsequently interpolated across all 80 imaging time-points (Supplemental Video 10). The resulting 2D masks were extruded along the left-right axis to encompass the complete gut volume within the z-stack imaging boundaries.

### J. Image analysis

All image processing and data analysis were performed with code written in Python using the NumPy [62], SciPy [63], scikit-image [64], and pandas [65] libraries. All code is available at https://github.com/rplab/Immune_Cell_Motion_Analysis_2025.

#### 1. Preprocessing

Each 2D slice of a 3D image volume (z-stack) was downsampled by a factor of 4 in both x and y directions to reduce computation time. Raw image stacks were then drift-corrected using phase correlation between bright field images from consecutive time points. For gut imaging, 4-5 overlapping fields of view along the anterior-posterior axis were acquired. Stage positions saved in the acquisition metadata were extracted from configuration files and represent global coordinates of the images which were then transformed to local pixel coordinates which was used to stitch different fields of view to generate complete gut volumes.

#### 2. Cell segmentation

Immune cells were segmented using a two-stage approach combining iterative thresholding with morphological Active Contour Without Edges (ACWE) [66, 67].

In brief, we applied iterative simple thresholding to get loose boundaries around objects. Initial thresholding used median + 2*σ*, where *σ* is the standard deviation of pixel intensities. Objects exceeding 10^5^ *µ*m^3^, considerably larger than the possible volume of a single cell, were iteratively re-thresholded at their local median + n*σ*, incrementing n until all objects fell below the volume limit or n reached 7. Objects smaller than a sub-cellular threshold of 10^3^ *µ*m^3^ were removed.

For each remaining object, we tightened/refined boundaries around the object using morphological ACWE with the thresholded region as the initial level set. The algorithm ran for 10 iterations with parameters: smoothing=1, *λ*_1_=1, *λ*_2_=4, with early stopping after 3 iterations of no change.

#### 3. Cell property analysis

Cell properties including volume, centroid, and mean intensity were extracted for each segmented object.

Surface area was calculated via a marching cubes algorithm on Gaussian-smoothed segmented objects with 5-pixel padding to prevent edge artifacts. Supplemental Video 11 shows the generated surfaces on the segmented macrophages. Sphericity quantified morphological complexity and was computed as:

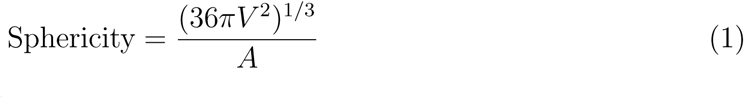

where *V* is cell volume and *A* is surface area. This metric equals 1 for perfect spheres and decreases with surface complexity, providing a quantitative measure of pseudopodial extension. As this metric is meaningful only between 0 and 1, the resulting values were clipped to this range.

Gut association was determined by spatial overlap with manually defined gut boundaries.

#### 4. Tracking and trajectory analysis

Segmented cells were tracked across time points using a bidirectional nearest-neighbor algorithm. For each consecutive time point pair, we computed Euclidean distances between all cell centroids in 3D space. A cell at time *t* was linked to its nearest neighbor at time *t* + 1 only if the reverse mapping (from *t* + 1 to *t*) also identified the original cell as the nearest neighbor. This bidirectional validation is a stringent criterion, such that tracks terminate when cells exit the imaging volume or when local crowding prevents unambiguous assignment; however, it also prevents spurious connections. The total number of tracked cells at each timepoint shows gradual changes without large fluctuations (Supplemental Figures 13 and 14), consistent with stable tracking over time. Trajectories were subdivided into “subtracks” at transitions above/below a displacement threshold of 0.5 × mean cell diameter, separating motile from non-motile phases. A patience parameter (n=3 timepoints) was applied to ensure subdivision only occurred after cells maintained the new motility state for 3 consecutive timepoints to prevent oversegmentation from transient behavior.

For each subtrack, we calculated:

- **Velocity components**: Frame-to-frame velocity components *V*_AP_, *V*_DV_, and *V*_LR_ along anterior-posterior (AP), dorsal-ventral (DV), and left-right (LR) axes
- **Velocity cosine similarity**: cos(*θ*) between consecutive velocity vectors
- **Spatial extent**: Diameter of smallest circumscribing sphere of the subtrack
- **Velocity anisotropy**: 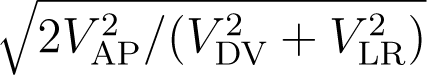

### K. Simulated random walk

To compare the observed spatial extent and velocity cosine similarity relations with what would arise from randomly directed motion, we generated simulated trajectories for each experimental track. For each track, we generated random walks with the same starting position, number of time points, and set of frame-to-frame step magnitudes, drawing step sizes at random and assigning directions drawn uniformly from the unit sphere. Positions were computed by cumulative summation from the original starting position. To replicate imaging, we applied simulated detection effects: Gaussian localization noise (*σ_xy_* = 0.21 *µ*m, *σ_z_* = 0.43 *µ*m) followed by quantization to the imaging pixel resolution (0.5 *µ*m in *xy*, 1.0 *µ*m in *z*). Simulated tracks were analyzed using the same pipeline applied to experimental data.

### L. Statistical analysis

Custom code written in Python was used for data analysis and plotting. All p values reported are from Mann–Whitney U tests, which are nonparametric and so do not assume normally distributed data. Unless otherwise noted, uncertainties are the 16th and 84th percentile of the distribution calculated from bootstrap subsampling.

## Supporting information

Supplemental Figures, Video Captions, and Dataset Descriptions

## ACKNOWLEDGMENTS

We thank Rose Sockol and the University of Oregon Zebrafish Facility staff for fish husbandry and care. We thank Catherine D. Robinson for performing gut dissections and bacterial plating, and Emily P. R. Avey for generating gut lumen masks for bacterial distribution analysis. This work was supported by the National Science Foundation under award 2310570 and the National Institutes of Health under award P01GM125576. The funders had no role in study design, data collection and analysis, decision to publish, or preparation. Figures 1A and 3A were created in Inkscape [68] using icons from BioRender.

## AUTHOR CONTRIBUTIONS

P. A. performed experiments and analyzed data. P. A. and R. P. designed the experimental and analysis methods and wrote the manuscript.

## DATA AVAILABILITY STATEMENT

Analyzed neutrophil and macrophage trajectories are available as Supplemental Information. Analysis code is publicly available at https://github.com/rplab/Immune_Cell_Motion_Analysis_2025.

