## Supplemental Figures, Video Captions, and Dataset Descriptions for "Characterizing trajectories of innate immune cells in larval zebrafish"

(Dated: January 13, 2026)

### I. SUPPLEMENTAL INFORMATION: CONTENTS

1. Supplemental Figures and Captions
2. Supplemental 3D Figure Captions
3. Supplemental Video Captions
4. Supplemental Dataset Descriptions

### II. SUPPLEMENTAL FIGURES AND CAPTIONS

*

†

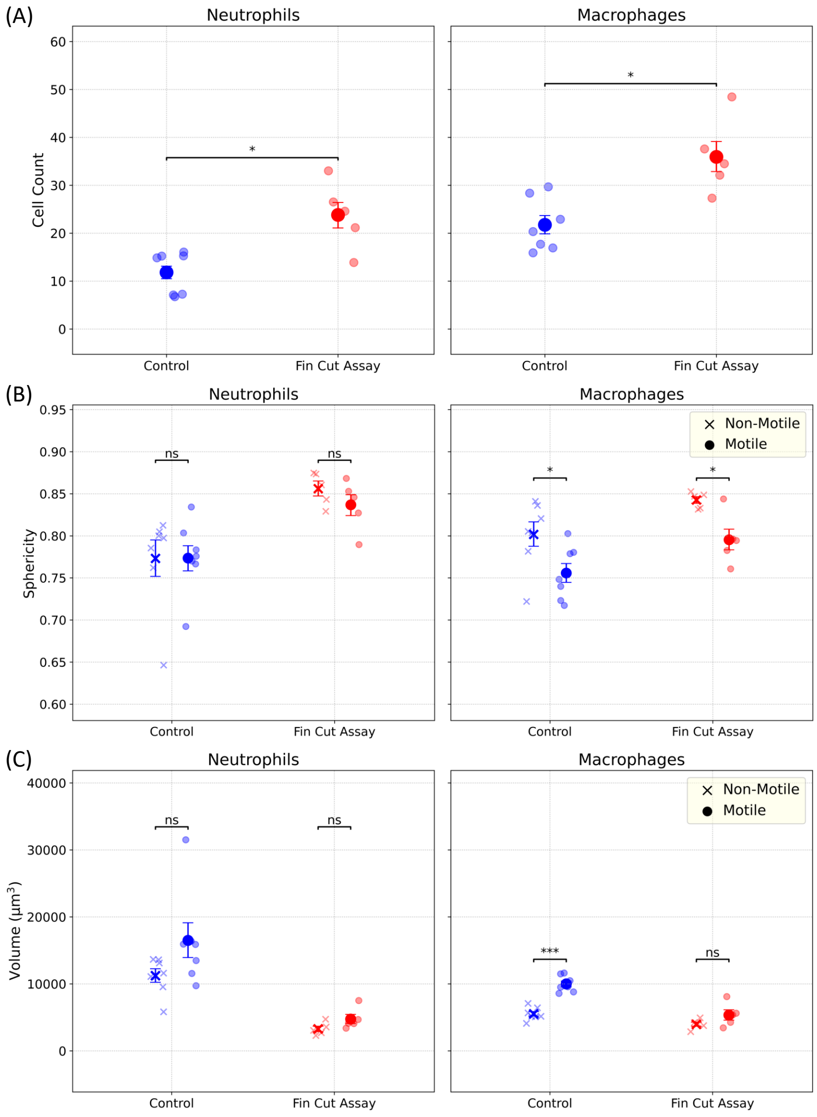


FIG. 1: Supplemental Fin Cut Assay: All panels show data for neutrophils (left) and macrophages (right). (A) Mean cell count in the field of view of imaging (approximately 800 $\mu$m from the posterior end of the caudal fin) over 1-5 hours post-amputation in control and fin cut conditions. Neutrophil numbers increased following fin amputation compared to controls, while macrophage counts remained unchanged between conditions. (B) Sphericity comparison between motile and non-motile subtracks across both experimental groups. Macrophages exhibit significantly higher sphericity during non-motile periods for both control and fin cut assay, indicating increased cellular rounding when stationary, while neutrophils show no morphological distinction between motility states. (C) Volume comparison between motile and non-motile subtracks in both experimental groups. Error bars represent uncertainties from bootstrapping. $n_{\text{Control}} = 5$, $n_{\text{Fin Cut Assay}} = 7$; total $N = 12$. ns : not significant, $* : p < 0.05$, $** : p < 0.01$, $*** : p < 0.001$.

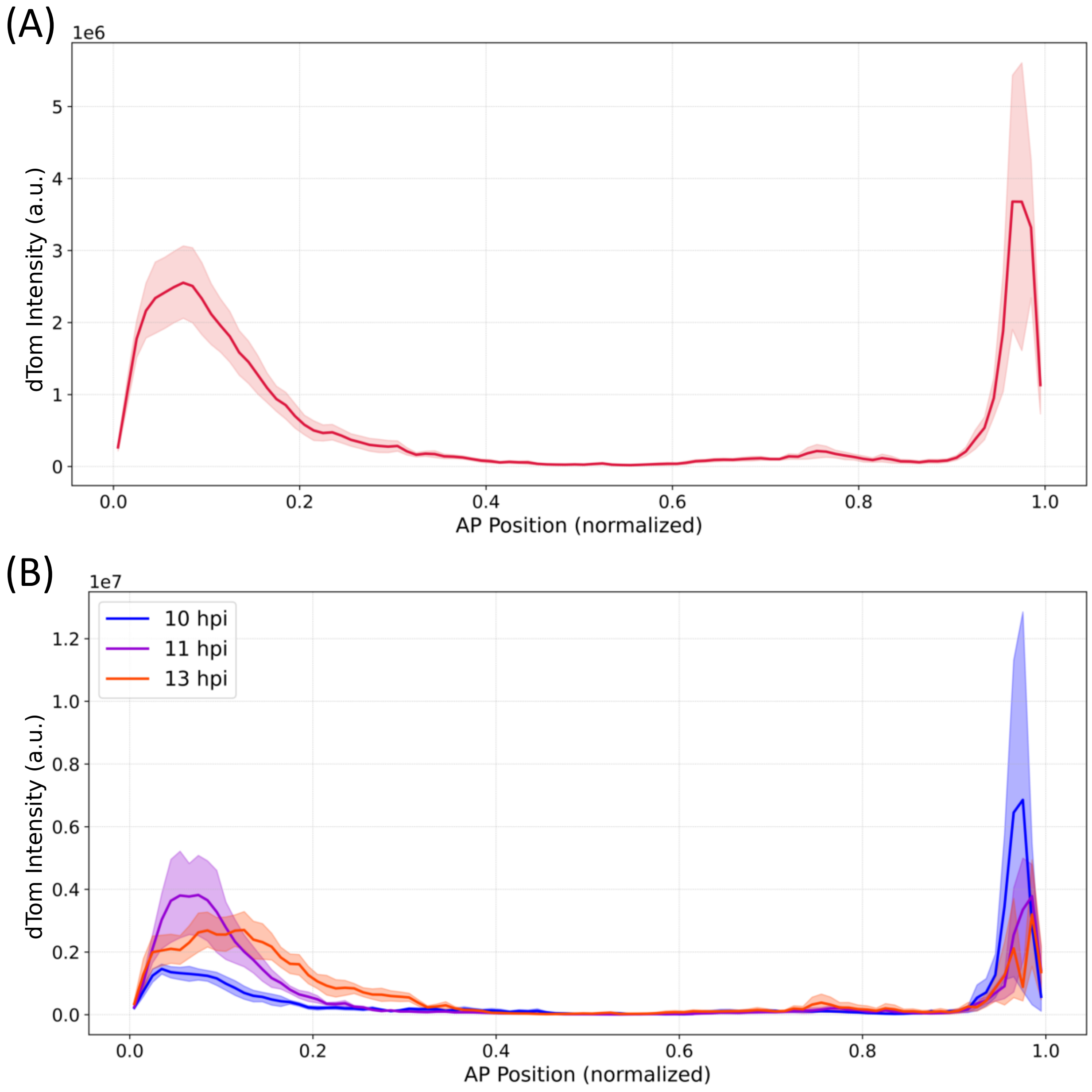


FIG. 2: Bacterial distribution and colonization during the imaging period. (A) Pooled median background-subtracted line profile of dTomato intensity along the anterior-posterior axis for $Vibrio^{WT}$-dTomato inoculated fish from 10-13 hpi. (B) Line profiles of dTomato intensity showing bacterial distribution at 10, 11, and 13 hpi. Bacterial fluorescence localizes primarily at the gut bulb (anterior) and the vent (posterior) at representative timepoints throughout the 9-13 hpi imaging window. Anterior-posterior position is normalized such that 0 and 1 represent the start and end of the gut, respectively. dTomato intensity is shown in arbitrary units (a.u.). Error bars represent uncertainties from bootstrapping. $n = 4$ per timepoint, total $N = 12$.

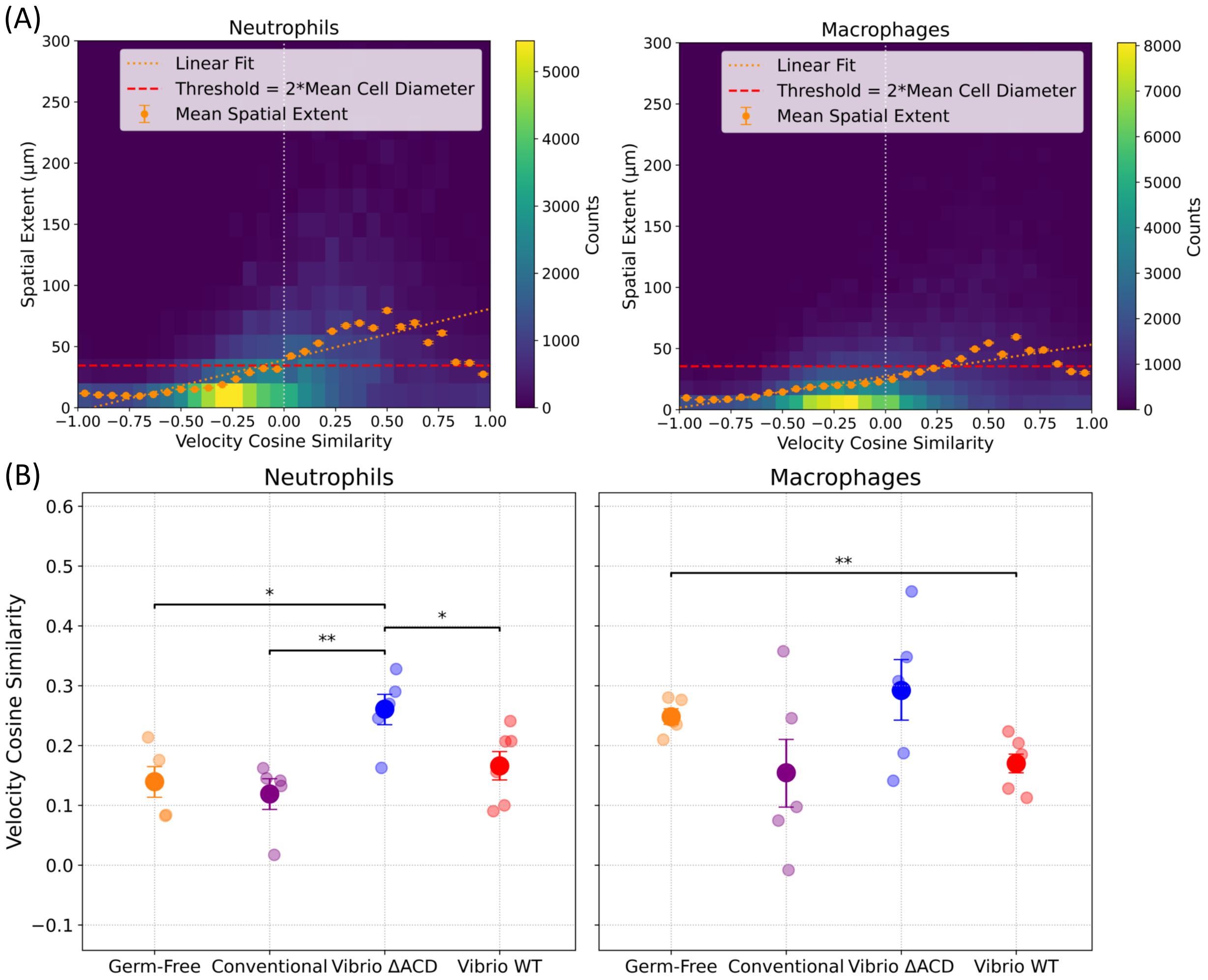


FIG. 3: Supplemental Gut Bacterial Association: All panels show data for neutrophils (left) and macrophages (right). (A) 2D histograms of spatial extent versus mean velocity cosine similarity for subtracks in the imaging region around the gut, pooled across control, conventional, and all *Vibrio*-associated fish. The horizontal red line shows the motility classification threshold of twice the mean cell diameter. Orange points show mean spatial extent binned by velocity cosine similarity; the orange dashed line is a linear regression fit to all individual data points. The y-intercept of the linear fit, i.e., spatial extent corresponding to zero velocity cosine similarity, is $38.98 \pm 0.23$ $\mu$m ($R^2 = 0.134$, $n = 1.40 \times 10^5$ datapoints) for neutrophils and $27.38 \pm 0.12$ $\mu$m ($R^2 = 0.120$, $n = 2.47 \times 10^5$) for macrophages. (B) Velocity Cosine Similarity for motile subtracks in control, conventional and *Vibrio*-associated fishes. $n_{\text{Germ Free}} = 5$, $n_{\text{Conventional}} = 5$, $n_{\text{Vibrio }\Delta\text{ACD}} = 5$, $n_{\text{Vibrio WT}} = 6$; total $N = 21$. $* : p < 0.05$, $** : p < 0.01$.

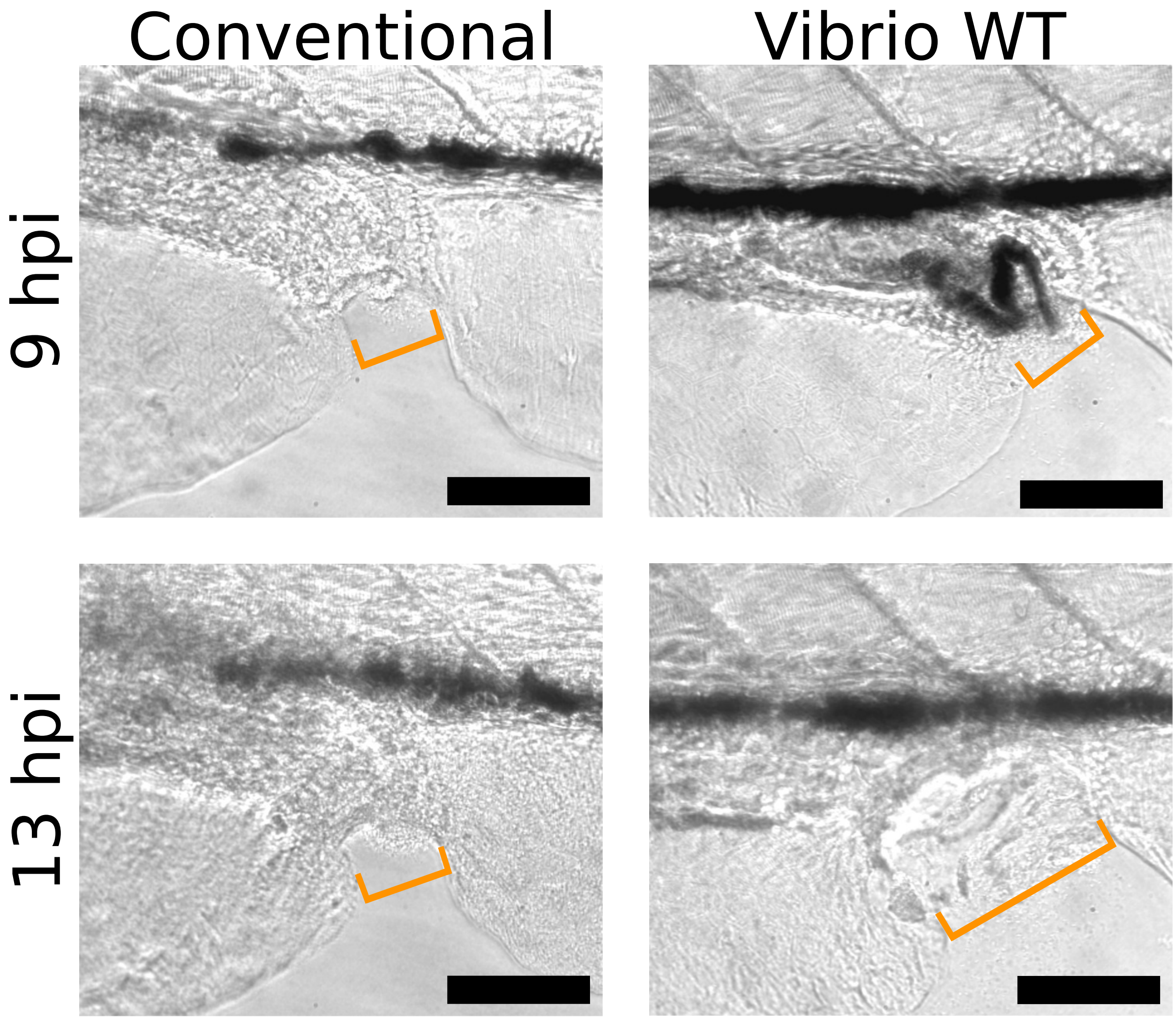


FIG. 4: Supplemental Gut Bacterial Association: Brightfield images showing vent widths for conventionally raised (left) and $Vibrio^{WT}$-inoculated fish (right) at the start and end of the imaging window. Orange lines indicate the vent width. All scale bars: 100 $\mu$m.

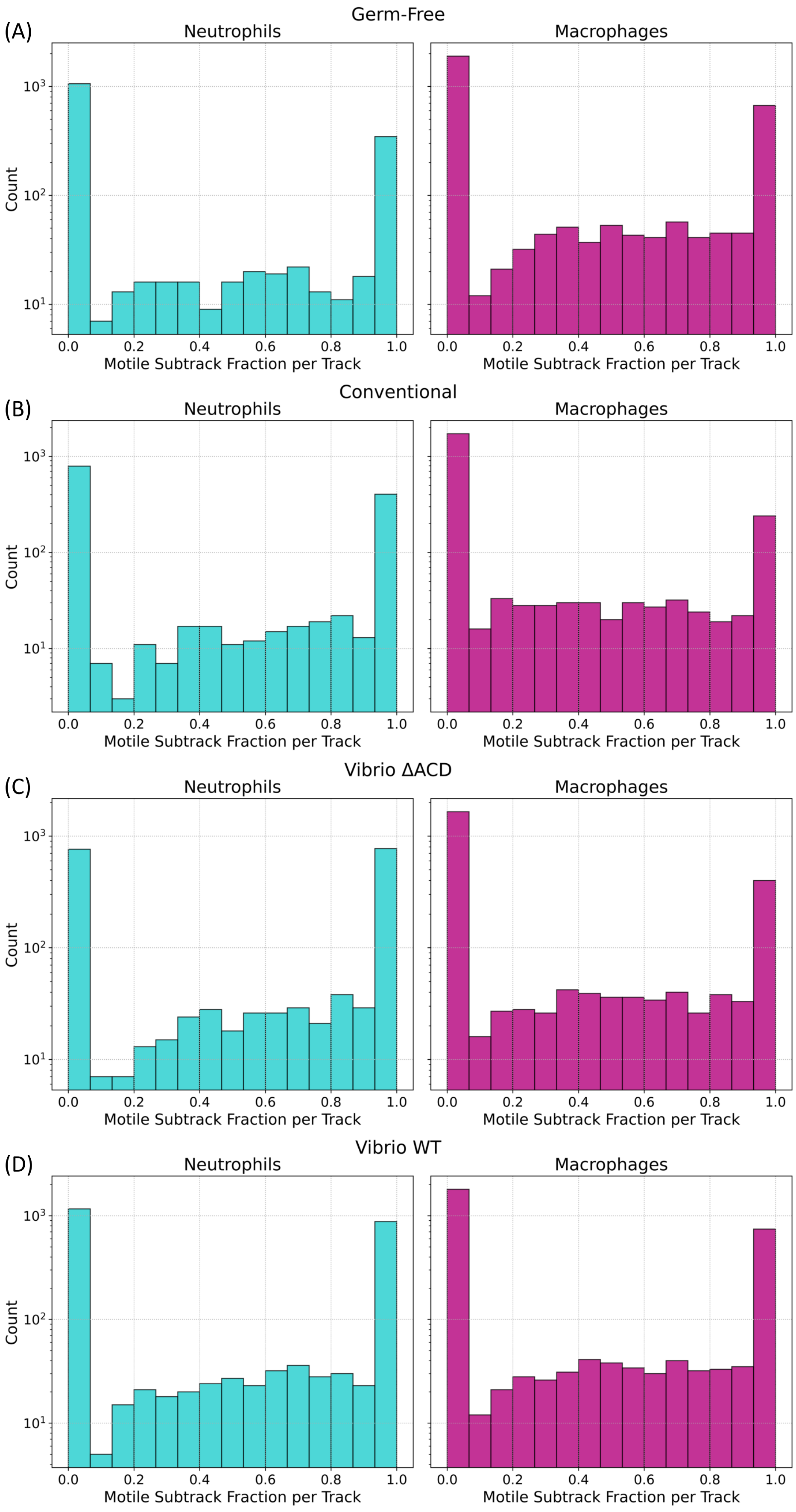

Germ-Free
(A)
Neutrophils
Macrophages
Count
Motile Subtrack Fraction per Track
Conventional
(B)
Neutrophils
Macrophages
Count
Motile Subtrack Fraction per Track
Vibrio ΔACD
(C)
Neutrophils
Macrophages
Count
Motile Subtrack Fraction per Track
Vibrio WT
(D)
Neutrophils
Macrophages
Count
Motile Subtrack Fraction per Track

FIG. 5: Supplemental Gut Bacterial Association: Histograms showing the fraction of each complete cell track exhibiting motile behavior for neutrophils (left) and macrophages (right), for (A) Germ-free (GF), (B) conventionally reared (Conventional), (C) $Vibrio^{\Delta ACD}$-inoculated, and (D) $Vibrio^{WT}$-inoculated fish. The distribution shows peaks at 0 and 1, indicating that immune cells generally adopt a consistent motile or non-motile phenotype over the 4-hour observation period, though a subset of cells exhibits intermediate behavior. Analysis included only tracks with minimum track lengths of 5 frames. Bin width = $0.0\overline{6}$. $n_{\text{Germ Free}} = 5$, $n_{\text{Conventional}} = 5$, $n_{\text{Vibrio }\Delta\text{ACD}} = 5$, $n_{\text{Vibrio WT}} = 6$; total $N = 21$.

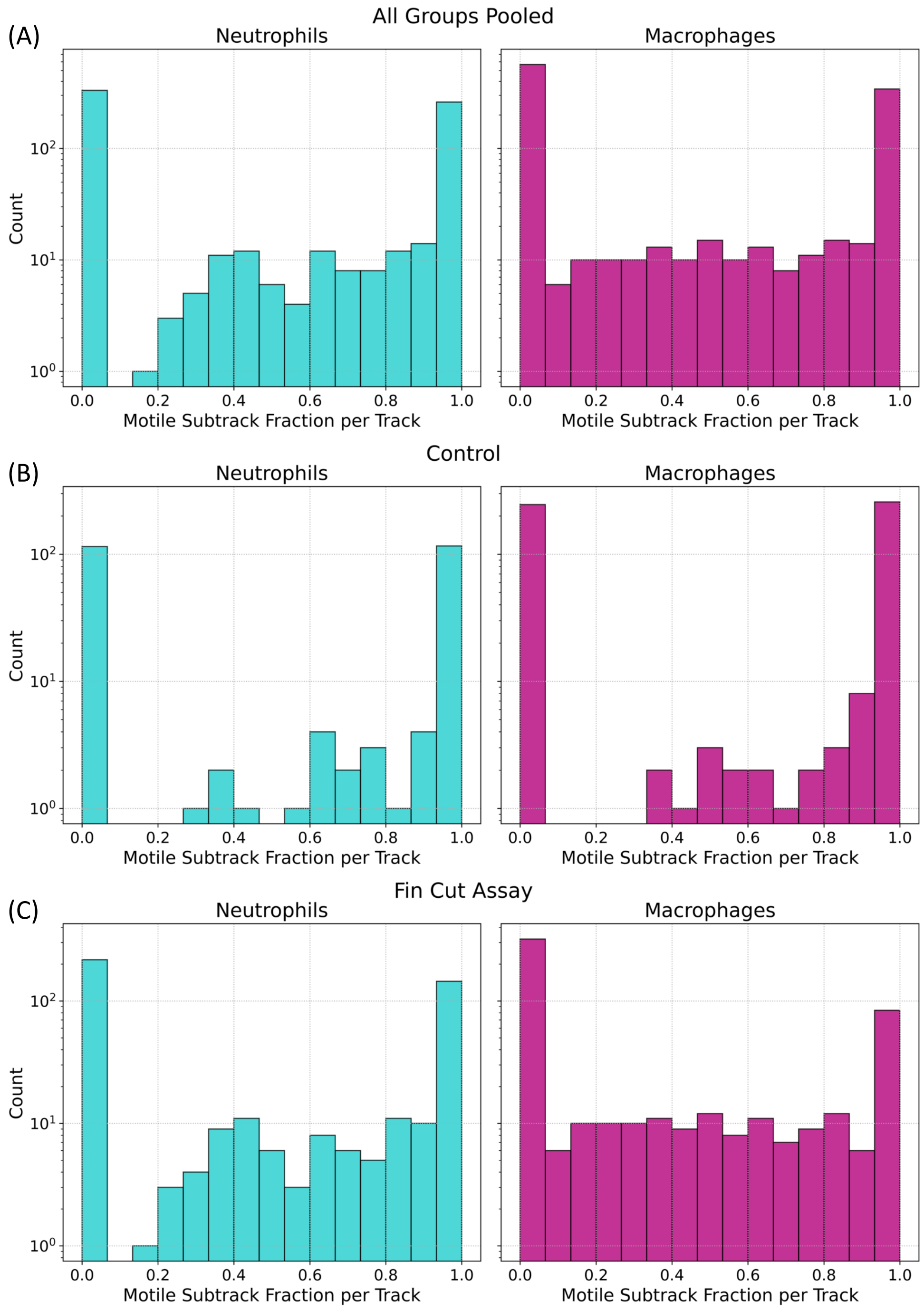


FIG. 6: Supplemental Fin Cut Assay: Histograms showing the fraction of each complete cell track exhibiting motile behavior for neutrophils (left) and macrophages (right), (A) pooled across all experimental groups, (B) only fin cut assay, and (C) only control (uninjured) group. The distribution shows peaks at 0 and 1, indicating that immune cells generally adopt a consistent motile or non-motile phenotype over the 4-hour observation period, though a subset of cells exhibits intermediate behavior. Analysis included only tracks with minimum track lengths of 5 frames. Bin width $= 0.0\overline{6}$. $n_{\text{Control}} = 5$, $n_{\text{Fin Cut Assay}} = 7$; total $N = 12$.

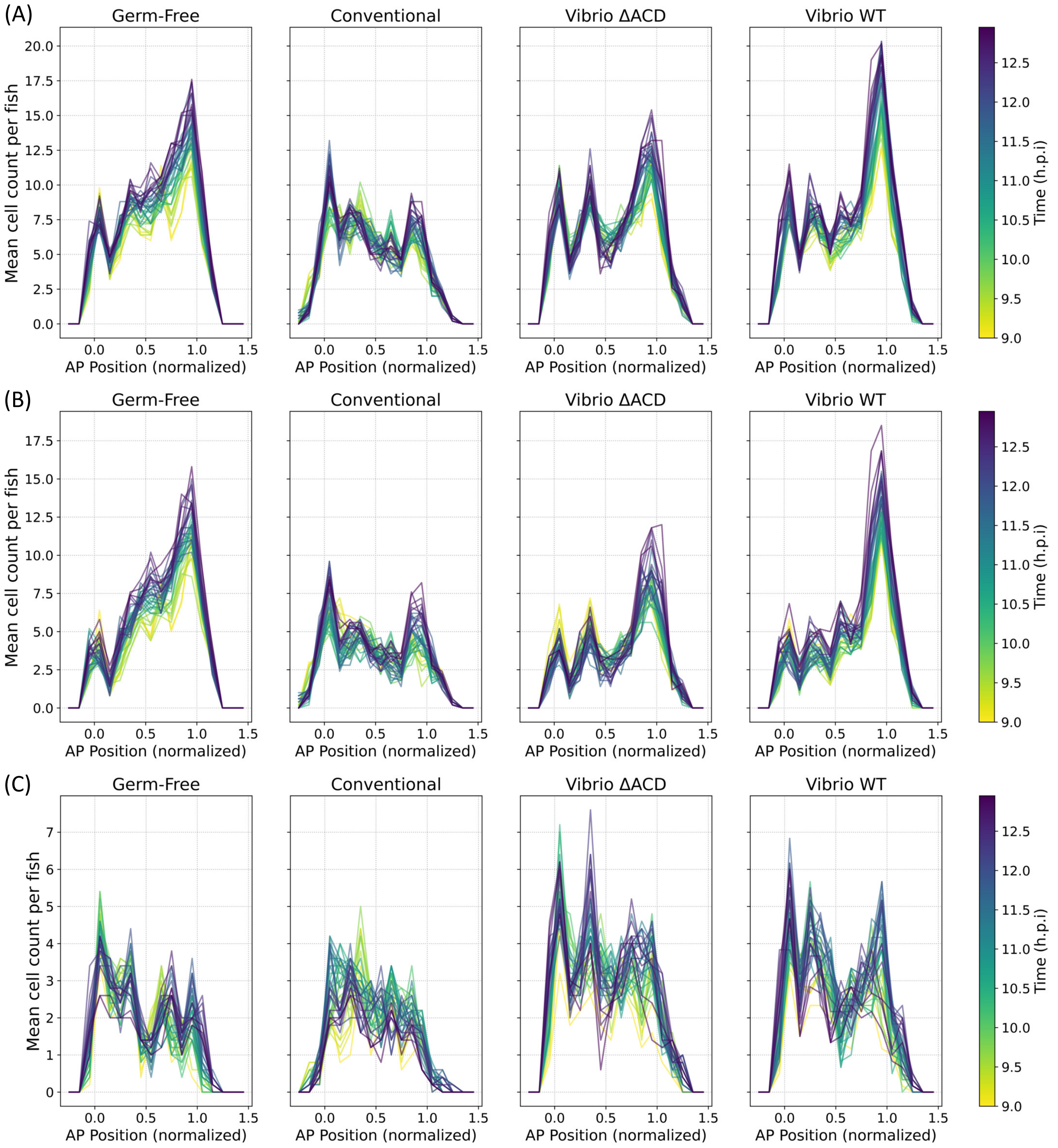


FIG. 7: Supplemental Gut Bacterial Association: Cell distribution along the anterior-posterior gut axis over time. Anterior-posterior position is normalized such that 0 and 1 represent the start and end of the gut, respectively. Color scale indicates time progression in hours post inoculation (h.p.i), where yellow represents early timepoints and dark purple represents late timepoints. (A) Total cell counts along the gut axis regardless of motility state. (B) Distribution of non-motile subtracks, showing increased cell accumulation near the vent region over time. (C) Distribution of motile subtracks. $n_{\text{Germ Free}} = 5$, $n_{\text{Conventional}} = 5$, $n_{\text{Vibrio }\Delta\text{ACD}} = 5$, $n_{\text{Vibrio WT}} = 6$; total $N = 21$.

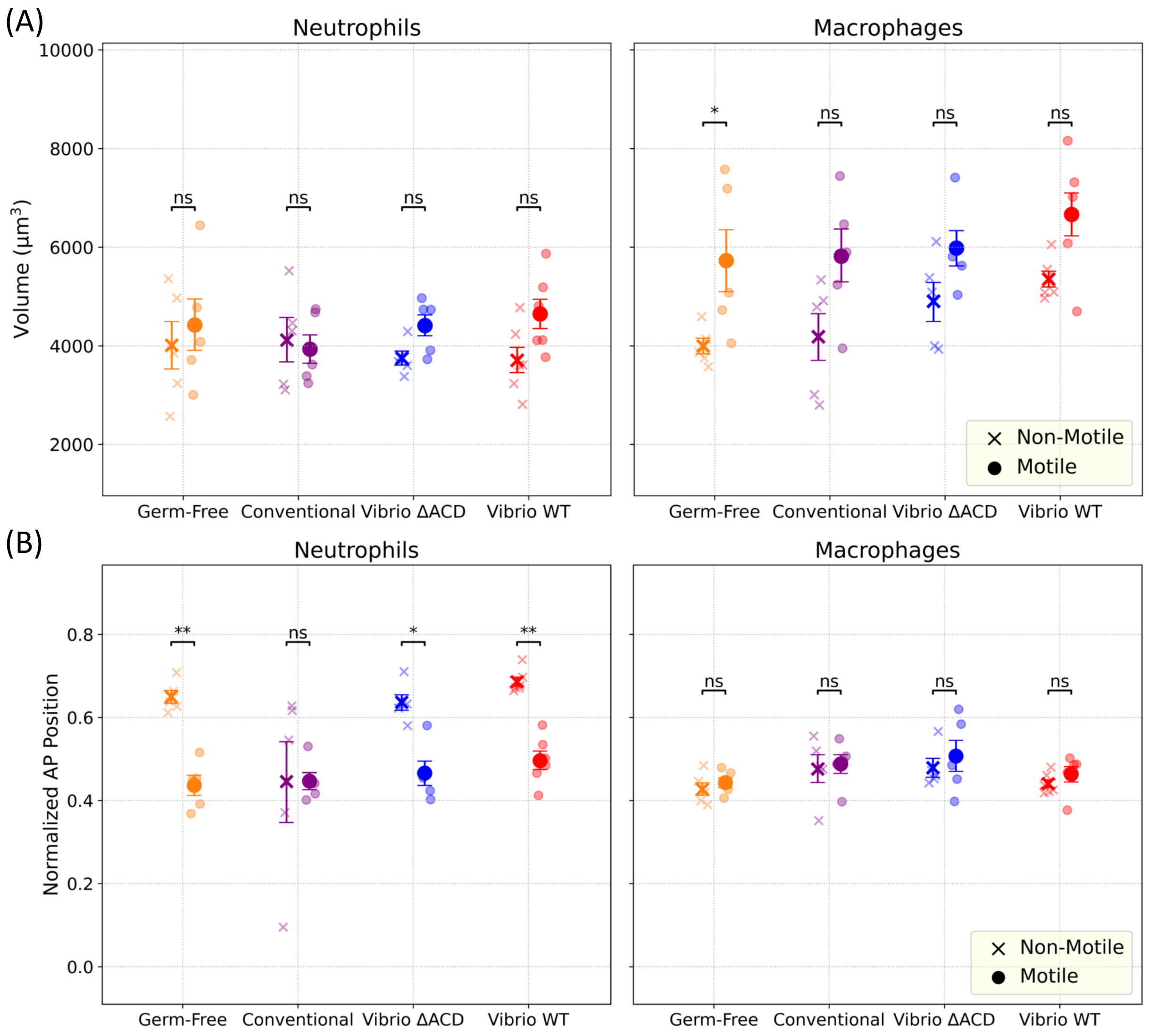


FIG. 8: Supplemental Gut Bacterial Association: (A) Volume comparison between motile and non-motile subtracks for neutrophils (left) and macrophages (right) across all experimental groups. Neither cell type shows significant differences between the cellular volumes. (B) Anterior-posterior position is normalized such that 0 and 1 represent the start and end of the gut, respectively. Non-motile neutrophils are generally more posterior located than motile neutrophils for all groups except the conventional. Macrophages show no significant difference in localization among the two motion types across all the groups, with average values around 0.5 implying they are equally distributed along the anterior-posterior axis. Error bars represent uncertainties from bootstrapping. $n_{\text{Germ Free}} = 5$, $n_{\text{Conventional}} = 5$, $n_{\text{Vibrio }\Delta\text{ACD}} = 5$, $n_{\text{Vibrio WT}} = 6$; total $N = 21$. ns : not significant, $* : p < 0.05$, $** : p < 0.01$, $*** : p < 0.001$.

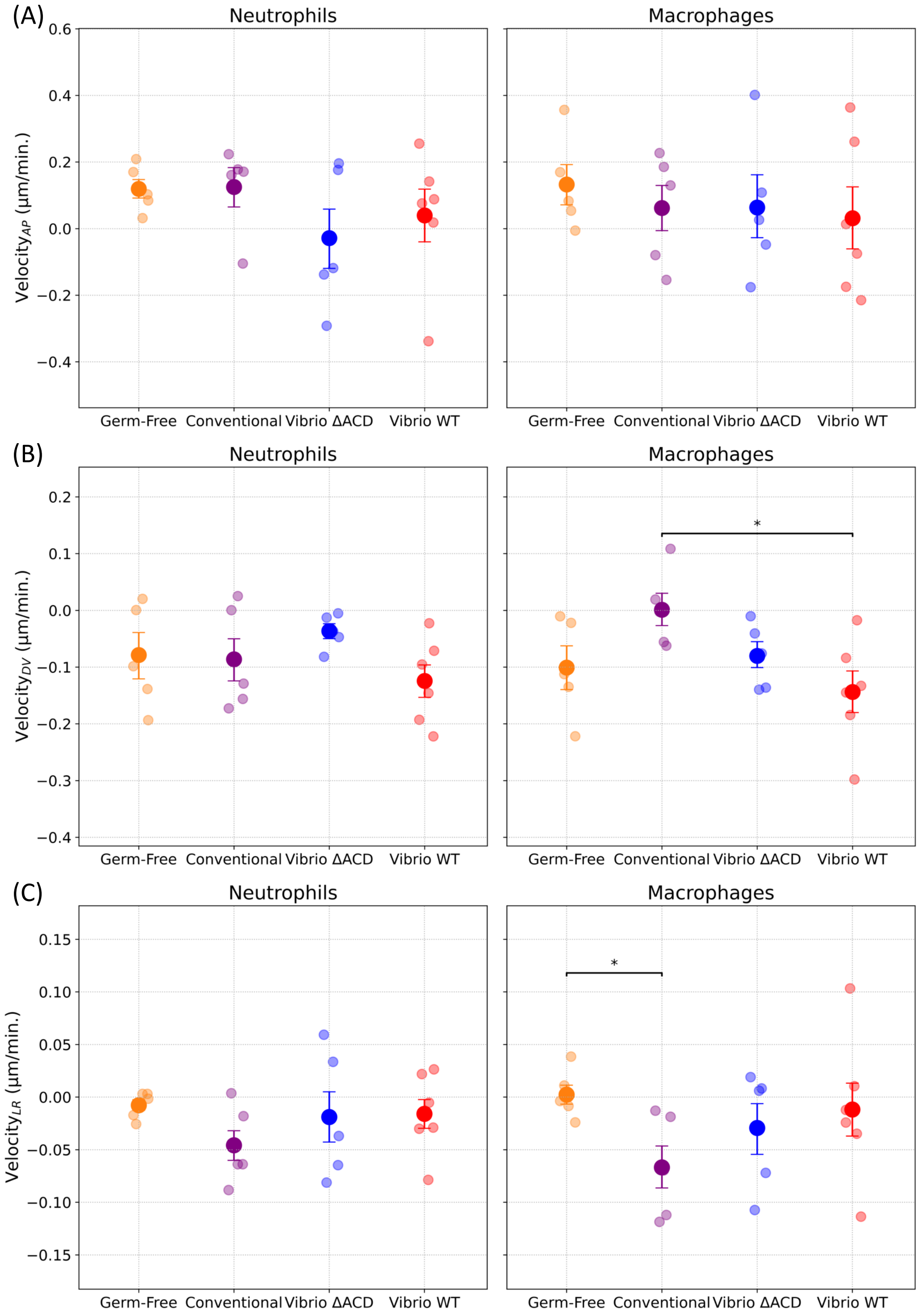


FIG. 9: Supplemental Gut Bacterial Association: The velocity component along the (A) anterior-posterior, (B) dorsal-ventral, and (C) left-right directions, calculated from frame-to-frame displacements, shows no significant differences between most experimental groups for either cell type. Error bars represent uncertainties from bootstrapping. $n_{\text{Germ Free}} = 5$, $n_{\text{Conventional}} = 5$, $n_{\text{Vibrio }\Delta\text{ACD}} = 5$, $n_{\text{Vibrio WT}} = 6$; total $N = 21$. $* : p < 0.05$.

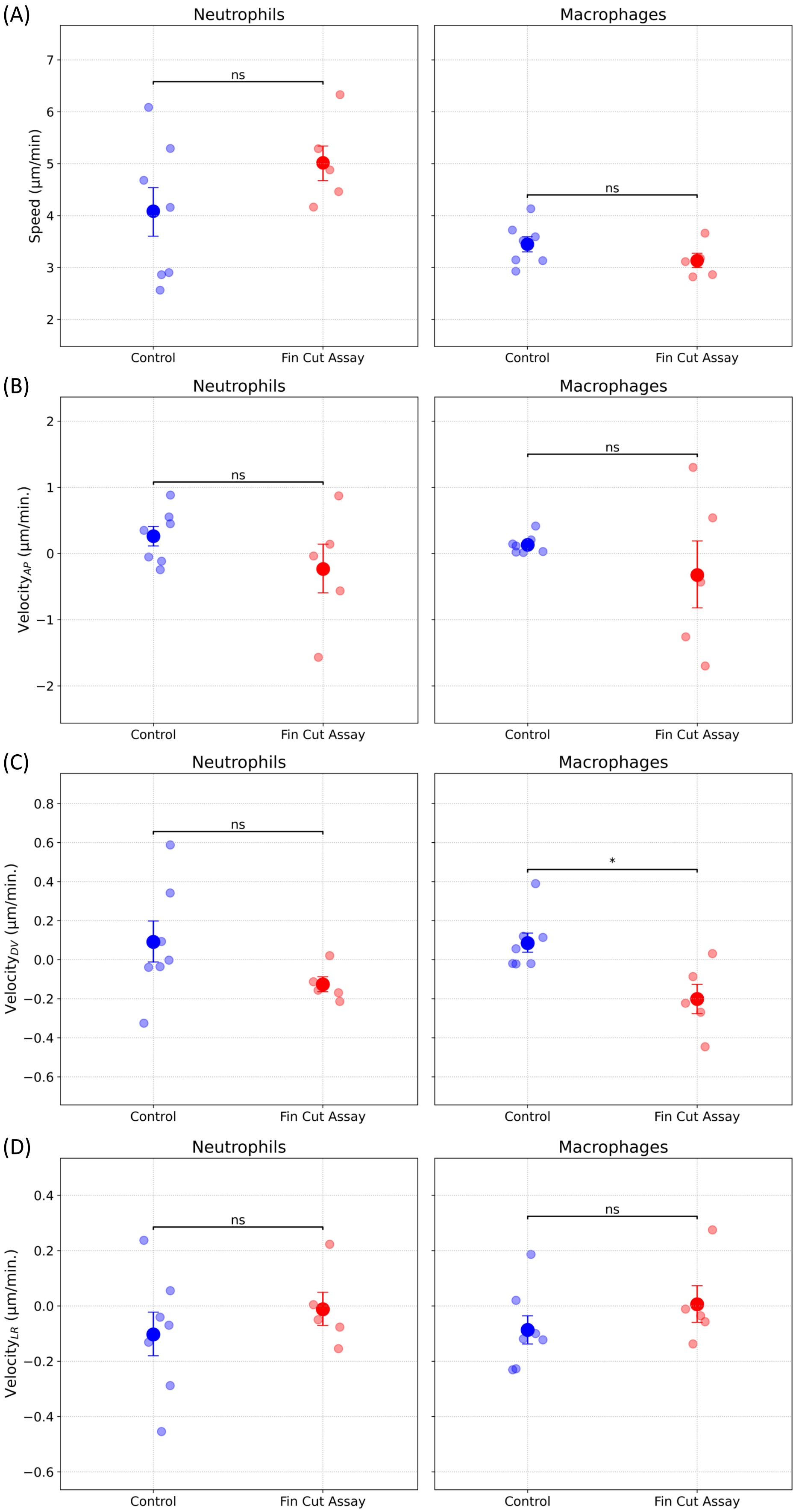

(A)
Neutrophils
Macrophages
Speed (μm/min)
ns
ns
Control
Fin Cut Assay
(B)
Neutrophils
Macrophages
Velocity$_{AP}$ (μm/min.)
ns
ns
Control
Fin Cut Assay
(C)
Neutrophils
Macrophages
Velocity$_{DV}$ (μm/min.)
ns
*
Control
Fin Cut Assay
(D)
Neutrophils
Macrophages
Velocity$_{LR}$ (μm/min.)
ns
ns
Control
Fin Cut Assay

FIG. 10: Supplemental Fin Cut Assay: The average speed (A) and the velocity component of motile subtracks along the (B) anterior-posterior, (C) dorsal-ventral, and (D) left-right directions, calculated from frame-to-frame displacements, shows no significant differences between experimental groups for either cell type. Error bars represent uncertainties from bootstrapping. $n_{\text{Control}} = 5$, $n_{\text{Fin Cut Assay}} = 7$; total $N = 12$. ns : not significant, $* : p < 0.05$, $** : p < 0.01$, $*** : p < 0.001$.

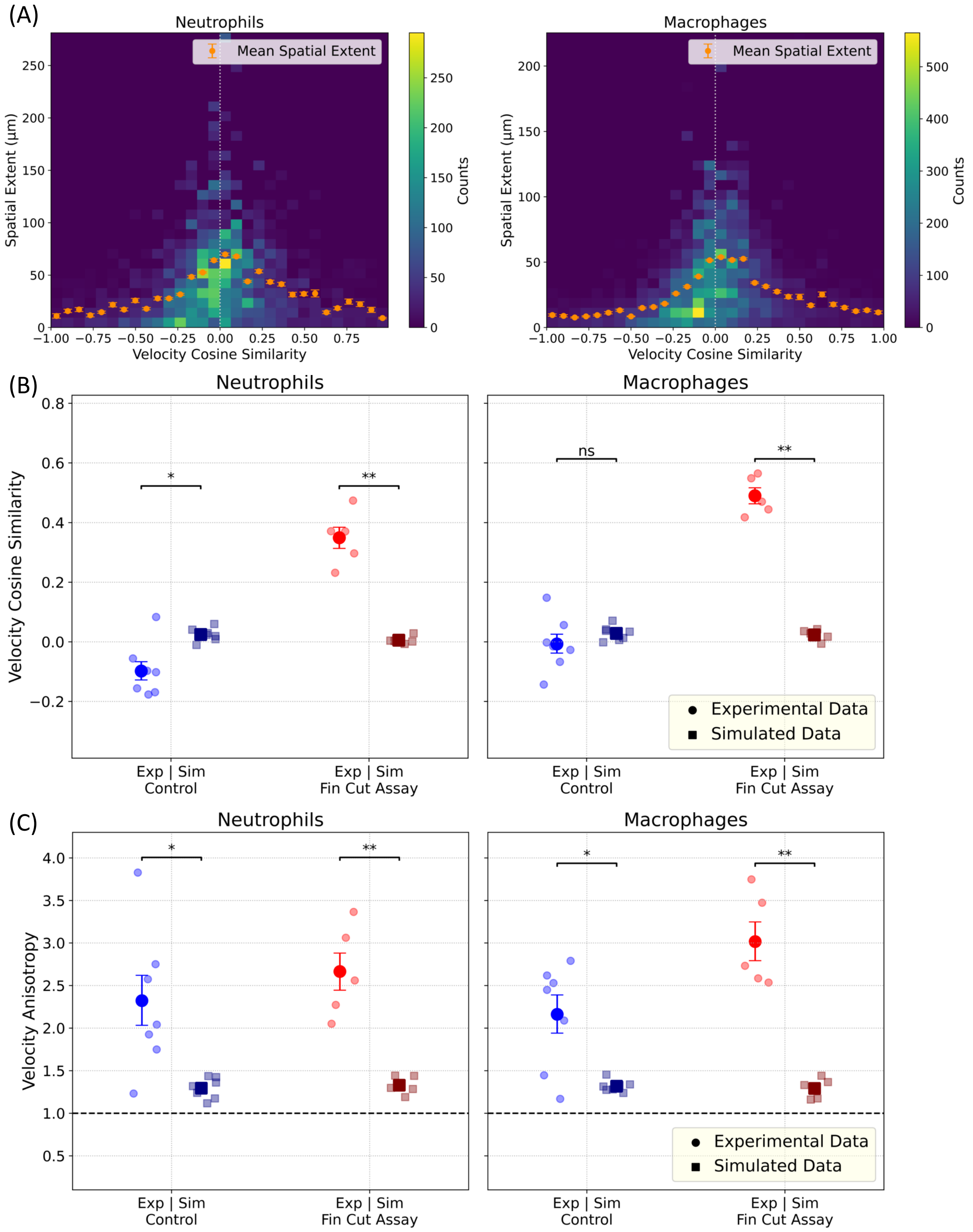
(A)
Neutrophils
Macrophages
Mean Spatial Extent
Spatial Extent (µm)
Velocity Cosine Similarity
Counts
(B)
Neutrophils
Macrophages
Velocity Cosine Similarity
ns
Experimental Data
Simulated Data
Exp | Sim
Control
Exp | Sim
Fin Cut Assay
(C)
Neutrophils
Macrophages
Velocity Anisotropy
Experimental Data
Simulated Data
Exp | Sim
Control
Exp | Sim
Fin Cut Assay

FIG. 11: Supplemental Simulated Fin Cut Assay: All panels show data for neutrophils (left) and macrophages (right). Simulated tracks were generated by randomizing step directions while preserving each track's step-size distribution and track length, with detector noise applied to match experimental imaging conditions (see Materials and Methods). (A) 2D histograms of spatial extent versus velocity cosine similarity for simulated subtracks, pooling control and fin-cut conditions. Orange points show mean spatial extent binned by velocity cosine similarity. Unlike experimental data (Figure 2A), simulated tracks show no linear correlation; velocity cosine similarity clusters almost symmetrically around zero regardless of spatial extent. (B) Velocity cosine similarity for simulated data clusters around zero, confirming random directionality. Experimental data show significantly higher directional persistence than simulated data for both cell types in the fin-cut condition. (C) Velocity anisotropy for motile subtracks. Dashed line indicates isotropy (anisotropy = 1). Experimental tracks exhibit significantly higher anisotropy than simulated tracks across all conditions. Error bars represent uncertainties from bootstrapping. ns : not significant, $* : p < 0.05$, $** : p < 0.01$.

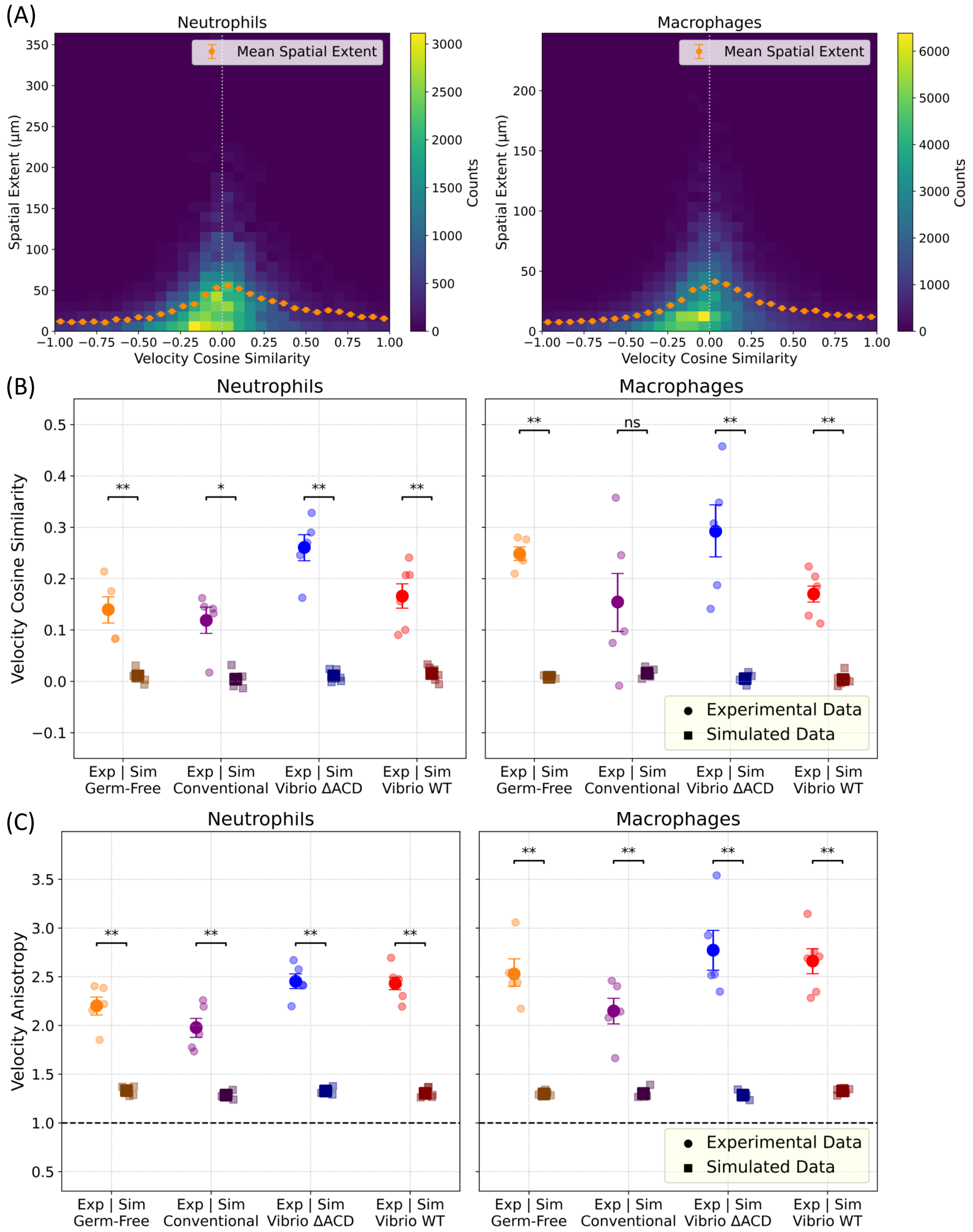
(A)
Neutrophils
Macrophages
Mean Spatial Extent
Spatial Extent (μm)
Velocity Cosine Similarity
Counts
(B)
Neutrophils
Macrophages
Velocity Cosine Similarity
**
*
ns
Experimental Data
Simulated Data
Exp | Sim
Germ-Free
Conventional
Vibrio ΔACD
Vibrio WT
(C)
Neutrophils
Macrophages
Velocity Anisotropy
**
Experimental Data
Simulated Data
Exp | Sim
Germ-Free
Conventional
Vibrio ΔACD
Vibrio WT

FIG. 12: Supplemental Simulated Gut Bacterial Association: All panels show data for neutrophils (left) and macrophages (right). Simulated tracks were generated by randomizing step directions while preserving each track's step-size distribution and track length, with detector noise applied to match experimental imaging conditions (see Materials and Methods). (A) 2D histograms of spatial extent versus velocity cosine similarity for simulated subtracks, pooling all conditions. Orange points show mean spatial extent binned by velocity cosine similarity. Unlike experimental data (Figure 3A), simulated tracks show no linear correlation; velocity cosine similarity clusters symmetrically around zero regardless of spatial extent. (B) Velocity cosine similarity for simulated data clusters around zero, confirming random directionality. Experimental data show significantly higher directional persistence than simulated data for both cell types across nearly all conditions. (C) Velocity anisotropy for motile subtracks. Dashed line indicates isotropy (anisotropy = 1). Experimental tracks exhibit significantly higher anisotropy than simulated tracks across all conditions for both cell types. Error bars represent uncertainties from bootstrapping. ns : not significant, $*: p < 0.05$, $**: p < 0.01$.

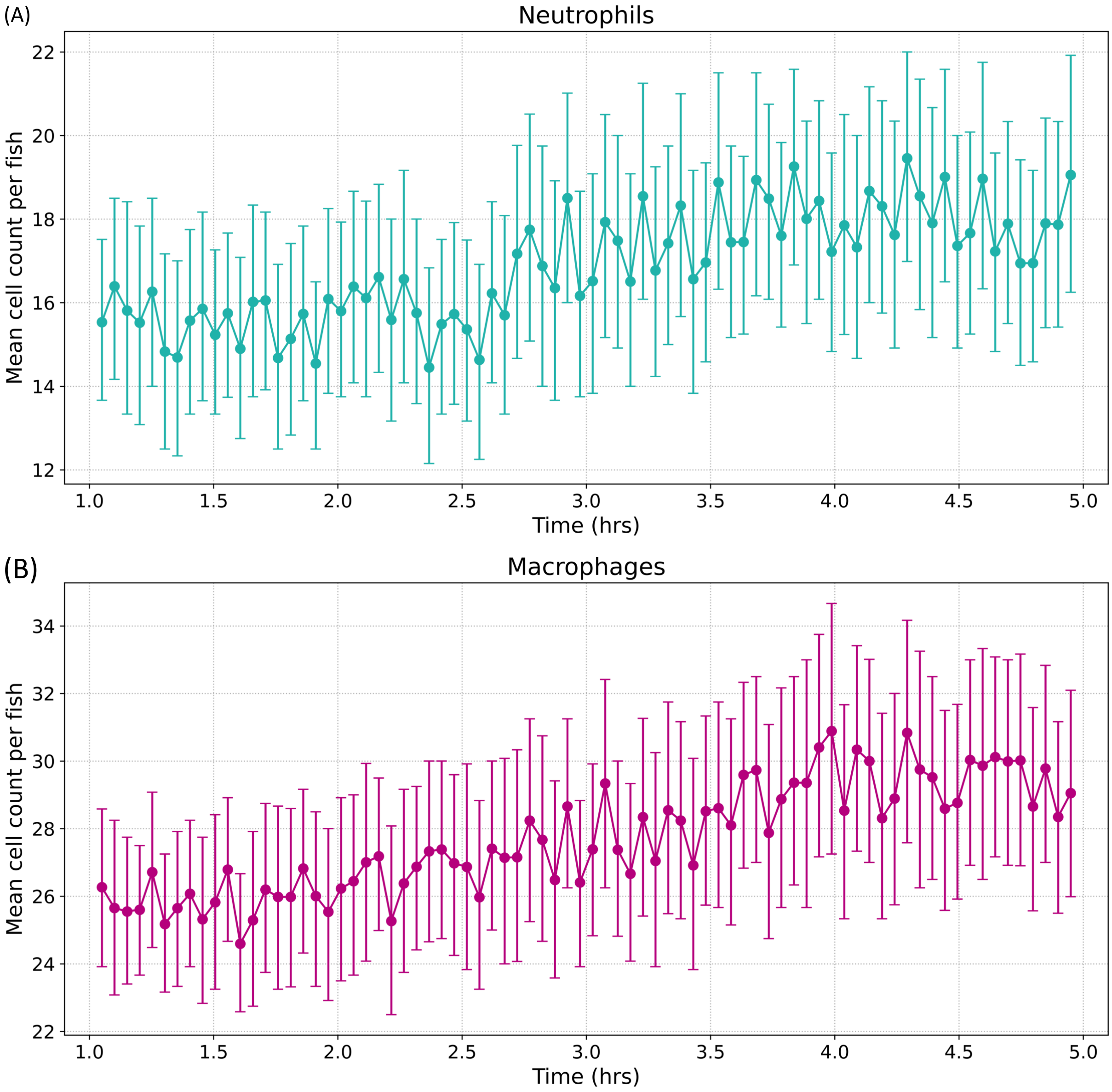


FIG. 13: Supplemental Fin Cut Assay: Mean cell count over time during fin cut assay experiments. (A) Neutrophils and (B) macrophages tracked in the caudal fin region from 1 to 5 hours post-amputation. Cell counts show gradual increases without large sudden fluctuations, indicating stable tracking of tissue-resident cells throughout the imaging period. Error bars represent standard error of the mean. Data pooled across all fin cut assay fish. $n_{\text{Control}} = 5$, $n_{\text{Fin Cut Assay}} = 7$; total $N = 12$.

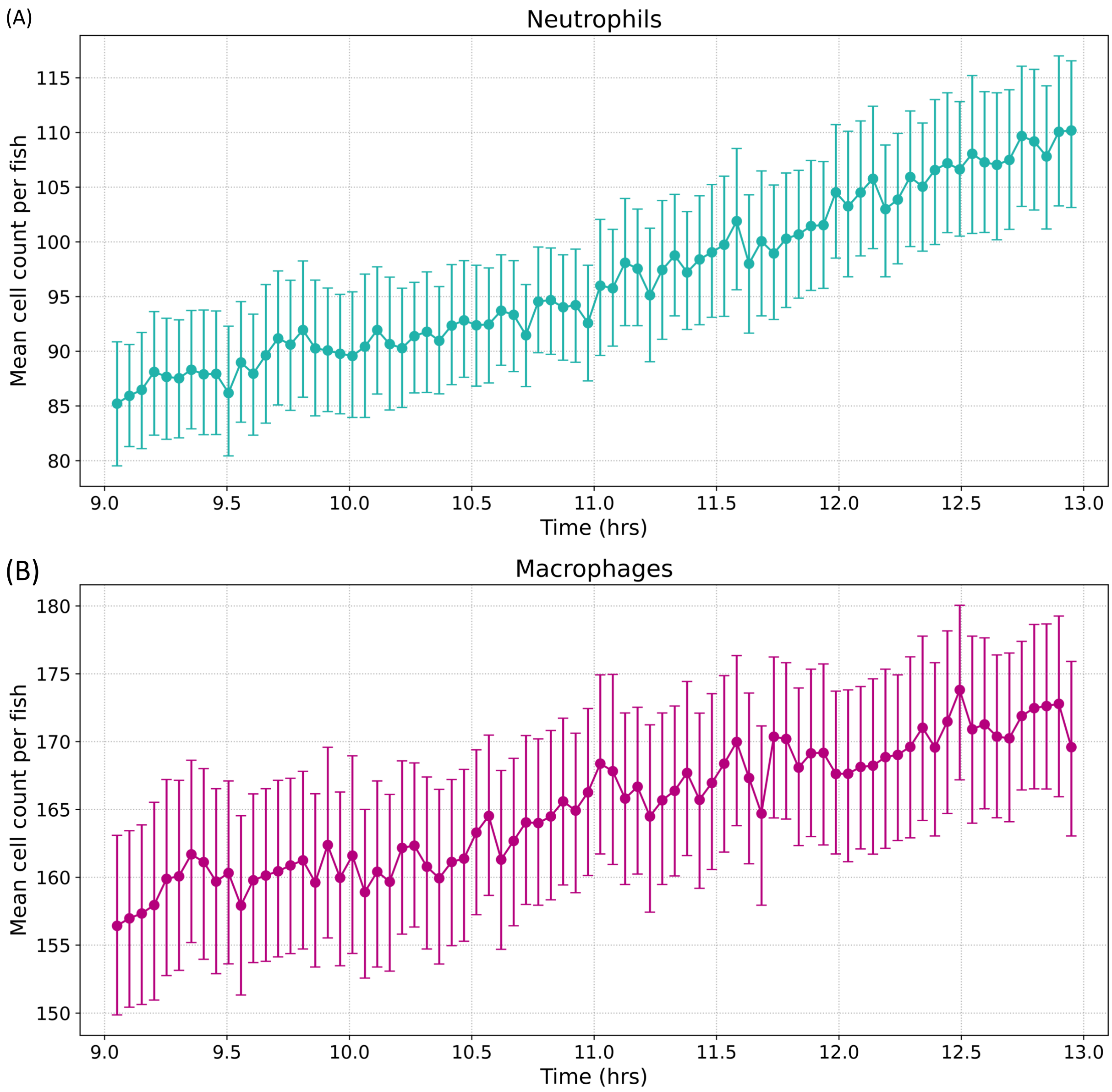


FIG. 14: Supplemental Gut Bacterial Association: Mean cell count over time during gut imaging experiments. (A) Neutrophils and (B) macrophages tracked in the gut region from 9 to 13 hours post-inoculation. Cell counts increase gradually without large sudden fluctuations, indicating stable tracking of tissue-resident cells throughout the imaging period. Error bars represent standard error of the mean. Data pooled across all experimental groups. $n_{\text{Germ Free}} = 5$, $n_{\text{Conventional}} = 5$, $n_{\text{Vibrio }\Delta\text{ACD}} = 5$, $n_{\text{Vibrio WT}} = 6$; total $N = 21$.

### III. SUPPLEMENTAL 3D FIGURE CAPTIONS

Supplemental 3D figures are available at Parthasarathy Lab Dropbox folder.

1. **Supplemental 3D Figure 1. 3D neutrophil trajectories colored by normalized time**: Interactive 3D visualization of neutrophil migration trajectories during fin cut assay from 1 to 5 hours post amputation. Track colors represent normalized temporal progression from track initiation (yellow) to termination (purple), with intermediate timepoints shown in gradient. AP: anterior-posterior; DV: dorsal-ventral; LR: left-right. Data corresponds to 3D representation of Figure 1C.

2. **Supplemental 3D Figure 2. 3D macrophage trajectories colored by normalized time**: Interactive 3D visualization of macrophage migration trajectories during fin cut assay from 1 to 5 hours post amputation. Track colors represent normalized temporal progression from track initiation (yellow) to termination (purple), with intermediate timepoints shown in gradient. AP: anterior-posterior; DV: dorsal-ventral; LR: left-right. Data corresponds to 3D representation of Figure 1D.

3. **Supplemental 3D Figure 3. 3D neutrophil trajectories colored by track identity**: Interactive 3D visualization of neutrophil migration trajectories during fin cut assay from 1 to 5 hours post amputation. Each unique track is assigned a distinct color based on track ID. Hover information displays timepoint and additional parameters for individual trajectories. AP: anterior-posterior; DV: dorsal-ventral; LR: left-right.

4. **Supplemental 3D Figure 4. 3D macrophage trajectories colored by track identity**: Interactive 3D visualization of macrophage migration trajectories during fin cut assay from 1 to 5 hours post amputation. Each unique track is assigned a distinct color based on track ID. Hover information displays timepoint and additional parameters for individual trajectories. AP: anterior-posterior; DV: dorsal-ventral; LR: left-right.

### IV. SUPPLEMENTAL VIDEO CAPTIONS

Supplemental videos are available at Parthasarathy Lab Dropbox folder.

1. **Video 1. Fin cut assay composite movie**: Composite image showing fin cut in gray, neutrophils in cyan and macrophages in magenta. Both immune cell types can be seen migrating to the fin amputation to the right of the figure from 1 to 5 hours post amputation. Scale bar = 100 $\mu$m.

2. **Video 2. Segmented neutrophils in maximum intensity projection responding to caudal fin transection**: Neutrophil response in fin cut assay showing 3D image stacks overlaid with segmented cells, unique colors show different segmented objects from 1 to 5 hours post amputation. Scale bar = 100 $\mu$m.

3. **Video 3. Z-scan through zebrafish gut showing bacterial distribution**: Volumetric z-scan along the left-right axis of the zebrafish gut at 13 hours post-inoculation with $Vibrio^{WT}$-dTomato. Image orientation: anterior-posterior (left to right), dorsal-ventral (top to bottom). Orange lines show the outline of the manually segmented gut lumen boundary at every 10th z-slice (see Methods and Materials). Scale bar = 100 $\mu$m.

4. **Video 4. Neutrophil trajectories on 4D image stacks**: Neutrophil dynamics across the entire gut field of view showing 3D image stacks with cells overlaid with pink dots showing centroids and unique colors showing different trajectories over 4 hours of imaging. Scale bar = 100 $\mu$m.

5. **Video 5. Macrophage trajectories on 4D image stacks**: Macrophage dynamics across the entire gut field of view showing 3D image stacks with cells overlaid with pink dots showing centroids and unique colors showing different trajectories over 4 hours of imaging. Scale bar = 100 $\mu$m.

6. **Video 6. 3D visualization of motile macrophage morphology**: Representative motile macrophage with rotating field of view demonstrating prominent pseudopodial extensions and reduced sphericity characteristic of the motile state. Scale bar = 20 $\mu$m.

7. **Video 7. Temporal dynamics of motile macrophage**: Representative motile macrophage in static view showing dynamic pseudopodial protrusions over time. Scale bar = 25 $\mu$m.

8. **Video 8. 3D visualization of non-motile macrophage morphology**: Representative non-motile macrophage with rotating field of view demonstrating rounded morphology and increased sphericity characteristic of the non-motile state. Scale bar = 10 $\mu$m.

9. **Video 9. Temporal dynamics of non-motile macrophage**: Representative non-motile macrophage in static view showing minimal morphological changes and absence of pseudopodial activity over time. Scale bar = 5 $\mu$m.

10. **Video 10. Neutrophil trajectories on maximum intensity projection with gut mask**: Maximum intensity projection of 3D image stack demonstrating manual gut segmentation and neutrophil dynamics across the entire gut field of view. Neutrophils are shown in cyan; the brightfield image is gray, overlaid with pink dots showing centroids and unique colors showing different trajectories over 4 hours of imaging. Scale bar = 100 $\mu$m.

11. **Video 11. 3D surface mesh visualization of gut bulb macrophages**: 3D surface mesh visualization of segmented macrophages in the gut bulb region, demonstrating the conversion from segmented objects to three-dimensional surface representations for morphological analysis. Scale bar = 50 $\mu$m.

### V. SUPPLEMENTAL DATASET DESCRIPTIONS

CSV files containing all tracked cell positions and other information is available at Parthasarathy Lab Dropbox folder. The experimental datasets are:

1. **gut_neutrophil_track_data_simplified.csv**: Trajectory and subtrack data for neutrophils from gut bacterial association experiments.

2. **gut_macrophage_track_data_simplified.csv**: Trajectory and subtrack data for macrophages from gut bacterial associations experiments.

3. **fin_cut_assay_neutrophil_track_data_simplified.csv**: Trajectory and subtrack data for neutrophils from fin cut assay experiments.

4. **fin_cut_assay_macrophage_track_data_simplified.csv**: Trajectory and subtrack data for macrophages from fin cut assay experiments.

The corresponding simulated datasets are available at Parthasarathy Lab Dropbox folder. The simulated datasets are:

1. **simulated_gut_neutrophil_track_data.csv**: Trajectory and subtrack data for neutrophils from gut bacterial association experiments.

2. **simulated_gut_macrophage_track_data.csv**: Trajectory and subtrack data for macrophages from gut bacterial associations experiments.

3. **simulated_fin_cut_assay_neutrophil_track_data.csv**: Trajectory and subtrack data for neutrophils from fin cut assay experiments.

4. **simulated_fin_cut_assay_macrophage_track_data.csv**: Trajectory and subtrack data for macrophages from fin cut assay experiments.

The columns in each CSV file are:

#### A. Identification and Temporal Columns

- **track_id**: Unique identifier for each complete cell trajectory
- **subtrack_id**: Unique identifier for trajectory subdivisions based on motility state transitions
- **time_point**: Frame number in the time series
- **slice**: Z, Y, X-slice position in the image stack according to numpy array slicing convention
- **fish_id**: Unique identifier for each individual zebrafish containing information about the experimental group.
- **dpf**: Days post fertilization at time of imaging

- **time_post_treatment_mins**: Minutes elapsed since treatment
- **time_post_treatment_hours**: Hours elapsed since treatment

#### B. Morphological Measurements

- **volume**: Cell volume in pixels$^3$.
- **volume_um**: Cell volume in $\mu$m$^3$.
- **user_surface_area_um**: Cell surface area in $\mu$m$^2$
- **sphericity**: Measure of how closely the cell shape approximates a sphere (0-1 scale).
- **holes**: Number of holes detected in the cell mesh produced by marching cubes. Any object with non-zero holes is not a closed object and discarded from further computations.
- **intensity_mean**: Average fluorescence intensity in the cell.

#### C. Spatial Position

- **pos**: the relative position of the image stack during acquisition. Final Image stacks are produced by stitching images from multiple positions.
- **x-dv-px**, **y-ap-px**, **z-lr-px**: Intensity-weighted cell centroid coordinates in pixels (dorsal-ventral, anterior-posterior, left-right directions defined as x, y, z axes respectively). These are global coordinates for the final stitched 3D image stack.
- **x-dv-um**, **y-ap-um**, **z-lr-um**: Intensity-weighted cell centroid coordinates in $\mu$m (dorsal-ventral, anterior-posterior, left-right directions defined as x, y, z axes respectively). These are global coordinates for the final stitched 3D image stack.
- **mask_synced_y_ap_px_normalized**: Normalized anterior-posterior position in pixels using hand-drawn masks.

#### D. Motion Parameters

All parameters are computed per subtrack unless otherwise noted.

- **speed_um-per-min**: Frame-to-frame cell speed in $\mu$m/min.
- **velocity_x_um-per-min**, **velocity_y_um-per-min**, **velocity_z_um-per-min**: Frame-to-frame velocity components along each axis in $\mu$m/min.
- **distance_um**: Frame-to-frame distance traveled in $\mu$m. Computed per track.
- **distance_px**: Frame-to-frame distance traveled in pixels. Computed per track.
- **velocity_cosine_similarity**: Measure of directional persistence computed as cosine of angle between consecutive velocity vectors.
- **spatial_extent_um**: Diameter of smallest circumscribing sphere of the subtrack in $\mu$m.
- **anisotropy_index_normalized**: Measure of trajectory directional bias along the anterior–posterior axis relative to the other two axis. Normalized such that isotropic motion yields a value of 1.

#### E. Motility Classification

- **motility**: Classification of motility states, either "motile" or "non-motile".
- **distance_px > mean_radius/2**: Boolean indicating if displacement exceeds half the mean cell radius.
- **spatial_extent > 2*mean_diameter**: Boolean indicating if spatial extent exceeds twice the mean cell diameter.

#### F. Track Characteristics

- **subtrack_length**: Number of timepoints in each subtrack.
- **track_length**: Total number of timepoints in the complete trajectory.

- **label**: Segmentation-derived identifier for individual cells within each image stack. Together with `timepoint` and `pos`, this forms a unique composite key that unambiguously identifies each segmented cell event in the entire dataset.

#### G. Anatomical Context for Gut Bacterial Association Experiments

- **in_gut**: Boolean indicating whether cell is located within the gut region.
- **gut_associated**: Binary classification of gut association. True if the cell trajectory intersects with the manually defined gut boundary at any timepoint within the subtrack.
- **vent_width_um**: Width of the vent in $\mu$m. Manually measured from bright-field images.

#### H. Scale Information

- **Pixel to micron conversion:** Anisotropic voxel size with 1 pixel = 0.65 $\mu$m in x-y directions and 1 pixel = 1.0 $\mu$m in the z direction.
- **Temporal resolution:** 3 minutes/frame.

#### I. Analysis Parameters

- **Motility threshold:** 0.5$\times$ mean cell diameter
- **Patience parameter:** 3 timepoints for subtrack subdivision
